# Deep inverse modeling reveals dynamic-dependent invariances in neural circuit mechanisms

**DOI:** 10.1101/2024.08.21.608969

**Authors:** Richard Gao, Michael Deistler, Auguste Schulz, Pedro J. Gonçalves, Jakob H. Macke

**Affiliations:** Machine Learning in Science, University of Tübingen and Tübingen AI Center, Tübingen, Germany; VIB-Neuroelectronics Research Flanders (NERF); imec, Belgium; Department Empirical Inference, Max Planck Institute for Intelligent Systems, Tübingen, Germany

## Abstract

Neural population dynamics are shaped by many cellular, synaptic, and network properties. Not only is it important to understand how coordinated changes in circuit parameters alter neural activity, but also when dynamics are unaffected by—or *invariant to*—such changes. Computational modeling has revealed invariances in single neurons and small circuits that are thought to reflect their robustness against variability and perturbations. However, generalizing these insights to larger circuits in cortex and other brain areas remains challenging. A key bottleneck lies in inverse modeling of neural circuits with spiking network models, i.e., identifying parameter configurations that quantitatively match dynamics observed in neural recordings. Here, we present *Automated Model Inference from Neural Dynamics (AutoMIND)* for efficient discovery of invariant circuit model configurations. AutoMIND leverages a mechanistic model with adaptive spiking neurons and clustered connectivity, which displays a rich variety of spatiotemporal dynamics. Probabilistic deep generative models—trained on network simulations only—then returns *many* parameter configurations consistent with a given target observation of neural activity. Applied to several datasets, AutoMIND discovers circuit models of synchronous network bursting in human brain organoids across early development, as well as models capturing complex frequency profiles of Neuropixels recordings in mouse hippocampus and cortex. In each case, we obtain hundreds of configurations that compose a (nonlinear) parameter subspace in which population dynamics remain unchanged. Surprisingly, global and local geometries of the invariant subspace are not fixed, but differ for different dynamics. Together, our results shed light on dynamic-dependent invariances of circuit parameters underlying diverse population dynamics, while demonstrating the flexibility of AutoMIND for inverse modeling of neural circuits.

## Introduction

Neural populations across the brain exhibit a wide spectrum of collective dynamics, including asynchronous states during alert behavior [1–3], fast and slow oscillations associated with various cognitive processes [4, 5], scale-free transients near criticality [6, 7], synchronous bursts in early brain development [8–10], and more. This repertoire of dynamics is shaped by a variety of circuit mechanisms at cellular, synaptic, and network scales, and in turn forms the computational substrate underlying flexible behavior and cognition [11–14]. Conversely, circuit abnormalities lead to pathological dynamics indicative of neurological, psychiatric, and developmental disorders [5, 15, 16]. Thus, an overarching goal in neuroscience aims to understand how changes in circuit properties—and the interaction thereof—influence the dynamics of neural populations.

On the flip side, an oft-neglected but critical question is: what changes in circuit properties *do not* impact population dynamics? Studies of single neurons and small, dedicated circuits suggest that neural systems exhibit so-called degeneracy [17, 18]. For instance, different compositions of ion channel conductances give rise to identical excitability behavior in neurons [19–21], while multiple cellular and synaptic configurations can maintain the same rhythmic activity in central pattern generators [22–24]. In other words, degeneracy implies that specific neuronal dynamics are *invariant* to certain changes in the parameters of those systems. What do these invariances look like, and why do they matter?

In the simplest case, individual parameters can be considered “sloppy” [25], and their changes largely inconsequential, in contrast to “stiff” parameters where even tiny perturbations can alter dynamics (Fig. 1a, top left). A more common scenario is that stiff and sloppy directions are not aligned with individual parameters, but *combinations* of parameters (Fig. 1a, middle). In such cases, each parameter can tolerate large changes, but must be precisely coordinated to preserve system behavior (e.g., connection probability vs. strength). In larger neural systems with many interacting variables, such as cortical circuits, the structure of invariance may be non-trivial and could even—in theory—depend on specific dynamical regimes (Fig. 1a, middle vs. bottom). Functionally, invariance to changes in many noisy biological components may provide robustness [19, 20, 22, 26, 27] and reflect homeostatic or other compensatory mechanisms against perturbations [18, 21, 23, 28–31]. Furthermore, invariances reveal how detailed, microscopic models can be abstracted to derive simpler, macroscopic models of emergent phenomena at relevant scales [32–34]. Thus, characterizing invariances is critical for understanding the organizing principles of neural circuits, and of biological systems in general, where degeneracy appears to be ubiquitous across scales [17, 35].

**Figure 1.**
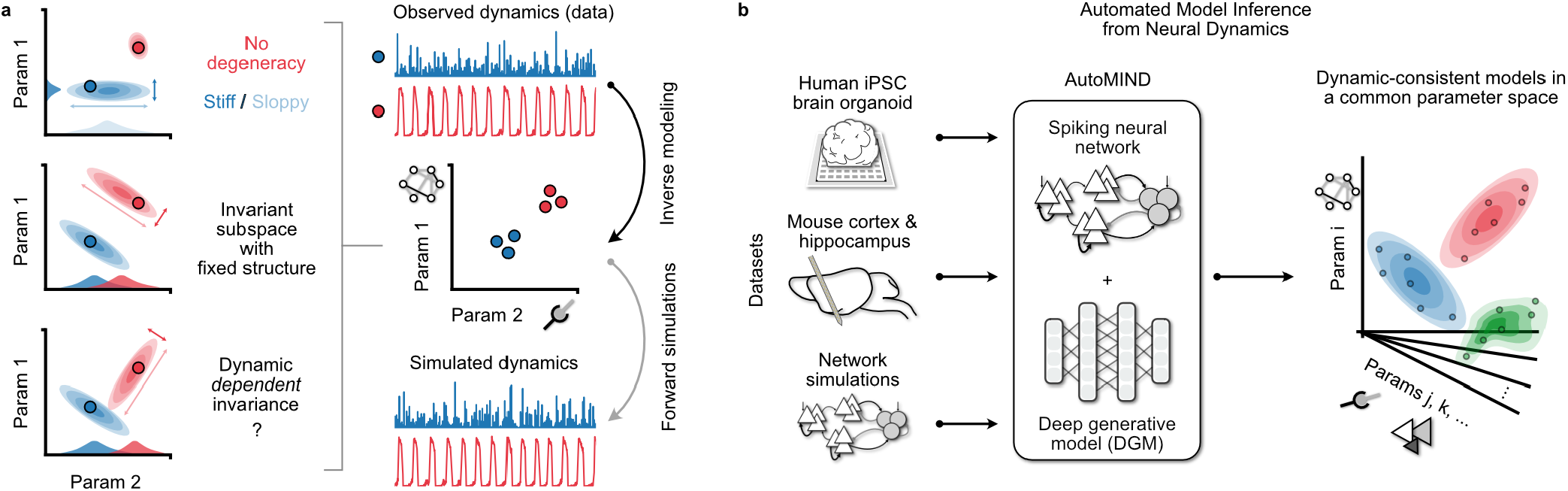
Schematic of structured parameter invariance revealed through inverse modeling. **a)** Inverse modeling seeks parameter configurations of mechanistic circuit models that are consistent with observations of neural dynamics. Discovered models may exhibit a variety of invariance structures (left column). Top: dynamic-consistent models may be unique (red), or have some parameters that are narrowly constrained (stiff) and others that are not (sloppy); Middle: stiff and sloppy directions may not be aligned with individual parameters, but combinations of parameters, defining a (potentially nonlinear) subspace on which dynamic is *invariant*. Bottom: the structure of the invariant subspace can, in theory, also depend on the target dynamic. **b)** AutoMIND leverages deep generative models to discover many dynamic-consistent parameter configurations of a common, flexible spiking network model architecture for different datasets.

Dissecting the sensitivity and invariance of neural dynamics to many circuit properties at once is often experimentally infeasible. To this end, network models of spiking neurons are invaluable, since their parameters encode mechanisms at different scales while balancing biophysical fidelity and tractability. In particular, by identifying parameter configurations that reproduce target observations of neural activity (i.e., *inverse modeling*), we can make inferences about unobserved quantities, test hypotheses with *in silico* interventions, and uncover potential invariances in the parameter space.

However, inverse modeling of neural circuits is challenging due to two mutually reinforcing problems: On the one hand, spiking network models are typically tailored to capture a specific type of dynamics or to test hypotheses about a few variables. As a result, they are often limited in the number of encoded mechanisms and free parameters, which also restricts their expressiveness to faithfully reproduce a broader range of dynamics. On the other hand, for larger, more expressive models with many free parameters, finding parameter configurations that are quantitatively consistent with observed dynamics becomes exponentially more difficult. Procedures for finding a single best-fitting configuration for just a handful of parameters are already computationally intensive, relying on brute-force search [22, 36] or iterative optimization routines [37–39], or require non-trivial properties of the model such as differentiability [40] or tractable likelihood functions [41]. An even bigger challenge lies in discovering *many different* configurations consistent with the same target observation—a necessity for characterizing invariances. Towards this, recent works leveraging (deep) generative models [42, 43] have shown improved computational efficiency [24, 44–48]. But due to the specificity and inflexibility of the mechanistic models used, they have been limited to applications targeting only simulated data or single experimental datasets. Altogether, these compounding issues prohibit investigations of circuit mechanisms underlying diverse neural population dynamics, and make it even harder to distill general principles of circuit invariances across disparate observations.

Here, we present *Automated Model Inference from Neural Dynamics* (AutoMIND), an inverse modeling framework for efficient discovery of mechanistic model parameter configurations consistent with observations of neural population dynamics (Fig. 1b). AutoMIND combines two ingredients to address the above challenges: First, we develop an expressive spiking neural network model that encodes as free parameters a variety of mechanisms at the cellular, synaptic, and network connectivity level, and exhibits a rich variety of temporal dynamics observed in experimental recordings. Second, leveraging probabilistic deep generative models and simulation-based inference [24, 49, 50], AutoMIND learns *solely* from a database of network simulations to enable efficient discovery of many parameter configurations that reproduce complex features of different experimental recordings.

Armed with these novel capabilities, we apply AutoMIND on three different datasets: longitudinal *in vitro* recordings from human brain organoid [51], *in vivo* Neuropixels recordings from mouse cortical and hippocampal areas [52], and diverse *in silico* model simulations with known parameter values. AutoMIND successfully discovers many parameter configurations of the same spiking network model that quantitatively reproduce target features of observed dynamics. Discovered models can further be perturbed to refine and test causal mechanistic hypotheses *in silico*. Finally, AutoMIND not only predicts individual parameter changes across developmental time, brain regions, and dynamical regimes, but reveals the existence of dynamic-dependent parameter invariances, suggesting complex, state-dependent interactions between cellular, synaptic, and network mechanisms underlying diverse neural population dynamics.

## Results

### Clustered spiking network model exhibits diverse population dynamics

The first ingredient to inverse modeling is a model with relevant mechanistic ingredients as free parameters, which is sufficiently expressive in reproducing key aspects of real experimental data. Here, we design a spiking neural network model that exhibits a wide range of population dynamics. The network is composed of 2000 adaptive exponential integrate-and-fire (AdExIF) neurons, which accurately models a range of single-neuron recordings [53, 54], split between excitatory (E) and inhibitory (I) populations (Fig. 2a). Neurons are connected through conductance-based synapses [55, 56], and network connectivity is sparse random but structured with excitatory neurons forming clusters [57]. Within-cluster excitatory connection probability and strength are amplified, while inhibitory neurons connect to all excitatory neurons with equal probability. These choices reflect the heterogeneous connectivity of a cortical circuit and neuronal adaptation, granting both long-timescale and spatially varying dynamics to population activity [58–61].

**Figure 2.**
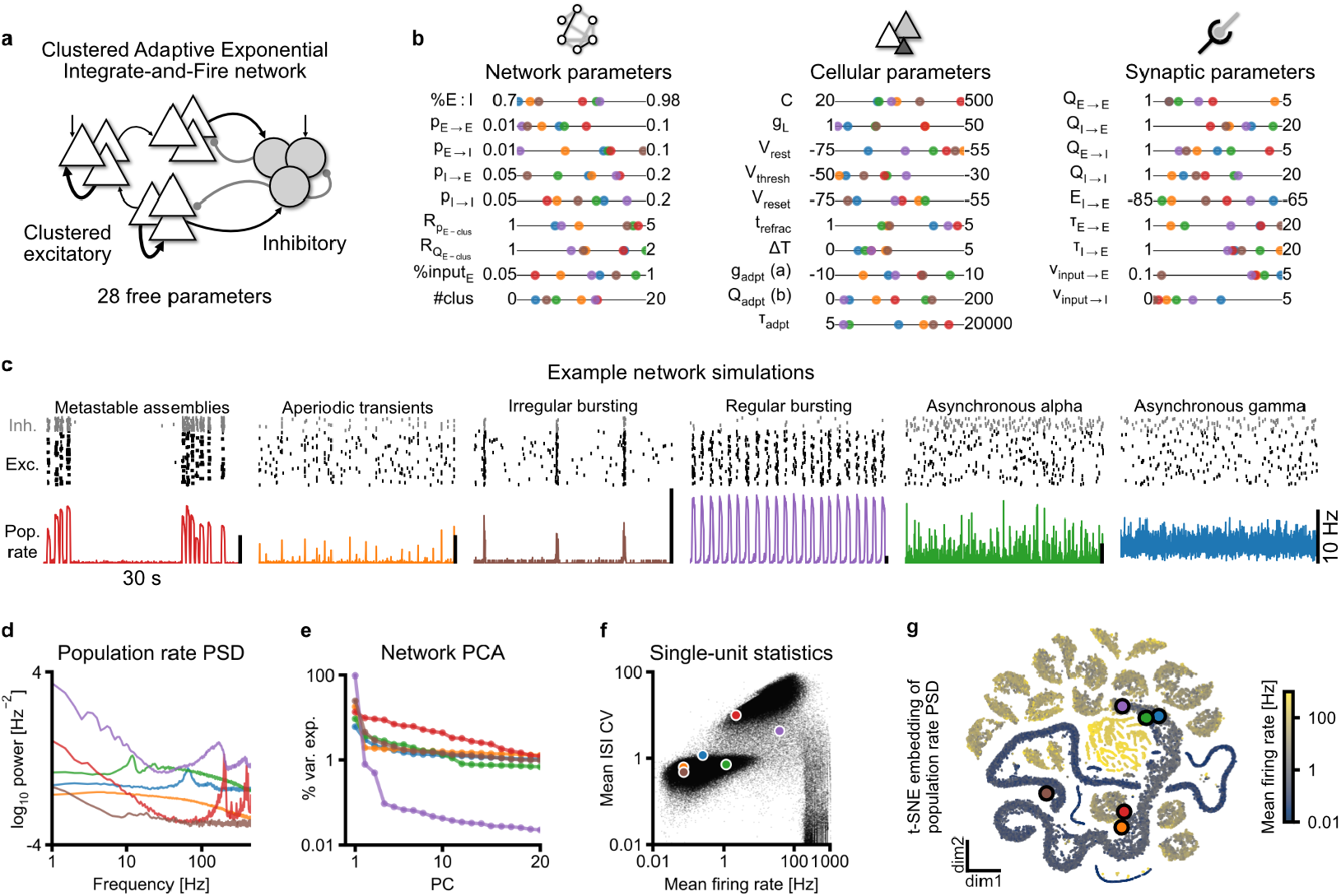
Clustered AdExIF model exhibits diverse and complex dynamical regimes. **a)** Schematic of clustered AdExIF network model architecture: excitatory populations can be clustered while an inhibitory population is shared; both receive external, excitatory Poisson input. **b)** 28 free parameters encode a variety of mechanisms at the network, cellular, and synaptic levels, such as connection probability and clusteredness, membrane and adaptation properties of excitatory neurons, synaptic strengths and time constants, and external input rates. Colors correspond to examples in **(c)**. Bounds denote the range of possible values (See Table 1 for details, including units). **c)** Example network simulations of parameter configurations in **(b**) showing a variety of possible dynamics, including assemblies, aperiodic and periodic transients and network bursts, as well as asynchronous oscillations of different frequencies. **d)** Power spectral densities (PSD) of population firing rates reveal periodic dynamics, such as 10Hz alpha (green), 65Hz gamma (blue), and other high-frequency oscillations (red, purple), as well as broadband 1/f-like behavior. **e)** Variance explained (log-scaled) across top 20 principal components (PCs); faster decay indicates fewer degrees of freedom and lower network dimensionality (e.g., purple). **f)** Networks exhibit a range of mean single-neuron firing rates and variability across 1 million random parameter configurations (black) with both gradual and abrupt transitions between dynamical states. ISI CV: inter-spike interval coefficient of variation. **g)** t-SNE embedding of 50,000 population rate PSDs highlights a continuous spectrum of low to moderate firing rate models distinct from islands of high firing rate networks. See Supplemental Fig. S2 for additional examples.

In total, the clustered-AdEx model has 28 free parameters (Fig. 2b, Table 1)—many more than typical spiking network models—which enable network simulations to display a larger range of possible dynamics while representing the circuit properties we want to infer from a given population recording. At the network scale, parameters such as the percentage of excitatory to inhibitory cells (%*E* :*I*) and connection probabilities (e.g., *p*_*E*→*E*_) can vary, as well as the number of clusters and within-cluster amplification. At the cellular level, membrane properties (*C, g*_*L*_, *V*_*rest*_, *V*_*thresh*_, etc.) and adaptation variables of excitatory neurons are free parameters. Lastly, synaptic properties, such as conductance at E and I synapses (e.g., *Q*_*E*→*E*_), their time constants, and external input rates are also free parameters. Thus, a vector of 28 numbers instantiates one model configuration (six examples in Fig. 2b), whose values are bounded between extremas aggregated from experimental data [62] or taken from previous modeling studies [63].

**Table 1.** Free parameters of the clustered AdEx network model, including their ranges (i.e., Uniform distribution support) and physical units.

| Parameter | Explanation | Range of values | Unit |
| --- | --- | --- | --- |
| *Network parameters* |  |  |  |
| $\%E:I$ | Percentage of excitatory neurons in the network | [70, 98] | % |
| $p_{E \rightarrow E}$ | Probability of excitatory to excitatory connection | [0.01, 0.1] | unitless |
| $p_{E \rightarrow I}$ | Probability of excitatory to inhibitory connection | [0.01, 0.1] | unitless |
| $p_{I \rightarrow E}$ | Probability of inhibitory to excitatory connection | [0.05, 0.2] | unitless |
| $p_{I \rightarrow I}$ | Probability of inhibitory to inhibitory connection | [0.05, 0.2] | unitless |
| $R_{pE-clus}$ | Amplification of within-cluster excitatory connection probability | [1, 5] | unitless |
| $R_{QE-clus}$ | Amplification of within-cluster excitatory synapse conductance | [1, 2] | unitless |
| $\%input_E$ | Percentage of excitatory neurons that receive input | [5, 100] | % |
| $\#clus$ | Number of clusters | [0, 20] | unitless |
| *Cellular parameters for excitatory neurons* |  |  |  |
| $C$ | Membrane capacitance | [20, 500] | $pF$ |
| $g_L$ | Leak conductance | [1, 50] | $nS$ |
| $V_{rest}$ | Resting potential | [-75, -55] | $mV$ |
| $V_{thresh}$ | Threshold potential | [-50, -30] | $mV$ |
| $V_{reset}$ | Reset potential | [-75, -55] | $mV$ |
| $t_{refrac}$ | Refractory period | [1, 5] | $ms$ |
| $\Delta_T$ | AP slope factor | [0, 5] | $mV$ |
| $g_{adpt}(a)$ | Subthreshold adaptation coupling | [-10, 10] | $nS$ |
| $Q_{adpt}(b)$ | Spike-triggered adaptation quantal current (b) | [0, 200] | $pA$ |
| $\tau_{adpt}$ | Adaptation time constant | [5, 20000] | $ms$ |
| *Synapses onto excitatory neurons* |  |  |  |
| $E_{I \rightarrow E}$ | Inhibitory to excitatory synapse reversal potential | [-85, -65] | $mV$ |
| $Q_{E \rightarrow E}$ | Excitatory to excitatory synaptic release conductance | [1, 5] | $nS$ |
| $Q_{I \rightarrow E}$ | Inhibitory to excitatory synaptic release conductance | [1, 20] | $nS$ |
| $\tau_{E \rightarrow E}$ | Excitatory to excitatory time constant | [1, 20] | $ms$ |
| $\tau_{I \rightarrow E}$ | Inhibitory to excitatory time constant | [1, 20] | $ms$ |
| *Synapses onto inhibitory neurons* |  |  |  |
| $Q_{E \rightarrow I}$ | Excitatory to inhibitory synaptic release conductance | [1, 5] | $nS$ |
| $Q_{I \rightarrow I}$ | Inhibitory to inhibitory synaptic release conductance | [1, 20] | $nS$ |
| *External input* |  |  |  |
| $\nu_{input \rightarrow E}$ | Poisson input rate to excitatory neurons | [0.1, 5] | $Hz$ |
| $\nu_{input \rightarrow I}$ | Poisson input rate to inhibitory neurons | [0, 5] | $Hz$ |

When endowed with such flexibility, the same network model can utilize the full parameter space to express a variety of realistic spatiotemporal dynamics (Fig. 2c), such as scale-free and metastable assembly dynamics, regular and irregular bursting with different periodicity, as well as asynchronous activity with or without oscillations. Simulations can be analyzed using standard methods in neuroscience: power spectral density (PSD) of the population firing rate uncovers characteristic features of neural recordings invisible from the raster, such as oscillatory peaks (e.g, 10 Hz alpha and 65 Hz-gamma), as well as broadband aperiodic, 1/f-like background [64] (Fig. 2d). Moreover, Principal Component Analysis (PCA) reveals varying network dimensionality (Fig. 2e), ranging from highly synchronous and low-dimensional, where the first principal component (PC1) explains almost 100 % of variance, to higher-dimensional dynamics with much slower eigenvalue decay [65]. This can be further summarized via measures such as participation ratio [66] (Supplemental Fig. S1a,b).

These examples illustrate the range of dynamics available to the clustered-AdEx network. To survey the broader parameter space, we simulated 1 million models instantiated by parameter configurations randomly drawn from the 28-dimensional uniform distribution defined in Table 1. After excluding invalid or pathological simulations (Supplemental Fig. S1c,d; see Methods for exclusion criteria), we visualize the ∼260,000 valid simulations by their average single-unit statistics (Fig. 2f). We observe an apparent separation of dynamical regimes reminiscent of the classical asynchronous (low firing rate and coefficient of variation, CV, near 1) vs. synchronous regular (high firing rate, low CV) distinction [2], with an intermediate irregular bursting regime (high CV).

Finally, embedding the broadband PSD of population firing rates in 2D with t-SNE [67], we observe further details in the discrete and continuous transitions between regimes of network dynamics (Fig. 2g). Coloring the embedding by non-spectral features (e.g., mean firing rate) reveals a continuous asynchronous regime, morphing into and surrounded by “islands” of synchronous, bursting dynamics (additional examples in Supplemental Fig. S2). Thus, our network model recapitulates a wide range of neural population dynamics at multiple spatial and temporal scales—many of which observed in experimental recordings—making it suitable for inverse modeling constrained by real data.

### Automated model inference from neural dynamics

With an expressive mechanistic model in hand, the second step of inverse modeling is to discover parameter configurations that can produce simulations quantitatively consistent with target features of experimental recordings. Towards this, we leverage simulation-based inference (SBI) [24, 49, 50, 68] with neural conditional density estimators (specifically, Normalizing Flows [69, 70]), which belong to the broader family of so-called deep generative models (DGM) [42, 43], i.e., probability distributions parameterized by deep artificial neural networks.

In brief, DGMs are *solely* trained on the simulation dataset above to approximate the relationship between parameter configurations and summary features of (simulated) network activity. Subsequently, when provided with a target observation, including real experimental data, we can query the trained DGM for parameter configurations that best reproduce it. In other words, given a forward model that maps parameter values to (simulated) neural activity, we seek to find a probabilistic inverse mapping from neural activity to parameter values *based on simulations alone*. In the language of Bayesian inference, the DGM approximates the posterior distribution over parameters conditioned on the neural data we want to explain.

Two features make this approach particularly suitable for inverse modeling of circuit dynamics. First, unlike optimization procedures that seek a single best-fitting parameter configuration, learning a probabilistic inverse mapping (i.e., posterior distribution) allows us to sample *multiple different* parameter configurations consistent with the same target observation. This capability is crucial for characterizing parameter invariances, since all model configurations from the posterior distribution conditioned on the same observation are, by definition, invariant. Second, once trained, the DGM can be efficiently sampled given any target observation, without retraining or Markov Chain Monte Carlo sampling. Relying on this *amortization* property, we extend Neural Posterior Estimation (NPE) [24, 71] by sampling around posterior modes (i.e., high-density regions) and subsequently filtering those samples based on their simulation quality (see Method for details). Together, the clustered-AdEx model and DGM-powered parameter discovery procedure constitute AutoMIND—Automated Model Inference from Neural Dynamics—which returns numerous model configurations whose simulations quantitatively match target observations of neural dynamics. We thus acquire *in silico* replicas of plausible real-world circuits underlying experimental recordings.

### Evolving circuit mechanisms of early neurodevelopment in human brain organoids

We first apply AutoMIND to uncover circuit changes during early neurodevelopment in human induced pluripotent stem cell-derived (iPSC) brain organoids [51]. Brain organoids were maintained on multi-electrode arrays, where they exhibit synchronous network bursts stereotypical of *in vitro* cultures [72, 73] and *in vivo* during early neurodevelopment [10]. Over the course of 40 weeks, network bursts become more frequent and variable, i.e., shorter inter-burst interval (IBI) and higher coefficient of variation (IBI CV), while burst duration goes through a non-monotonic trajectory, rapidly decreasing then slightly increasing (Fig. 3a,c).

**Figure 3.**
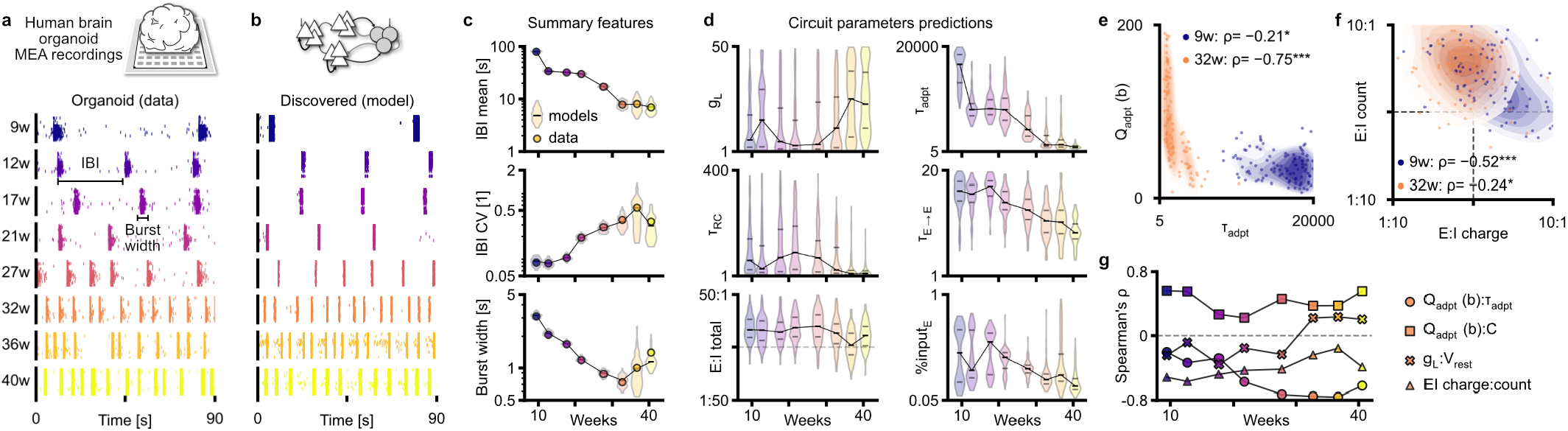
Circuit models of brain organoid development reveal evolving parameter distributions and invariances. **a)** Human iPSC-derived brain organoid recordings across 40 weeks, with evolving synchronized network dynamics (bursts) that decrease in inter-burst interval (IBI) and increase in stochasticity. **b)** Example of individual models consistent with observed recordings. **c)** Ensembles of 100 model configurations at each timepoint recapitulate target activity features of organoid data (lines connect median across models); CV: coefficient of variation. **d)** Distributions of a subset of model-derived circuit properties over developmental time; *g*_*L*_: leak conductance, *τ*_*RC*_ : membrane RC time constant, *E* :*I total* : ratio of total network synaptic excitation vs. inhibition, *τ*_*adpt*_ : adaptation time constant, *τ*_*E→E*_ : decay time constant of E-E synapse, %*input*_*E*_ : percentage of excitatory neurons receiving Poisson input. **e)** Increasingly negative correlation between adaptation current (*Q*_*adpt*_) and time constant in ensembles of models across two different timepoints (week 9 vs. 32). Spearman’s rho, */**/*** denote p<0.05, 0.01, 0.001 respectively. **f)** Decreasing correlation strength between E:I ratios of synaptic charge vs. synapse count. **g)** Evolving correlations between circuit properties in discovered model configurations over the course of early development. See Supplemental Fig. S4 for all parameters.

To dissect potential circuit mechanisms underlying these changes, we found candidate model configurations matching those dynamical features at each recorded timepoint: We first trained a DGM to learn the mapping between parameter configurations and burst summary features computed on the database of network simulations. Then, we queried the DGM for models consistent with organoid network burst features (Fig. 3b; additional examples in Supplemental Fig. S3). Critically, the probabilistic DGM returns many configurations for each recording over 40 weeks, all of which reproduce the target dynamical features observed *in vitro* (Fig. 3c, violin over 100 models).

With these dynamic-consistent models, we can examine how cellular and network parameters change over developmental time (Supplemental Fig. S4a). To highlight a few: we observe that leak conductance (*g*_*L*_) increases, and as a result, membrane time constant (*τ*_*RC*_) decreases, especially later in development, while networks are generally excitation-dominated until late (Fig. 3d, left columns). These observations corroborate existing experimental data of decreases in membrane resistance and membrane time constant in early postnatal rodents [74, 75] (Supplemental Fig. S4b), as well as the late emergence of GABAergic inhibition in brain organoids [51] and rodents [76].

Additionally, there are consistent decreases in neuronal adaptation time constant (*τ*_*adpt*_), excitatory synaptic time constant (*τ*_*E*→*E*_), and the percentage of input-receiving neurons, which represent spontaneously active neurons in the network (Fig. 3d, right columns). While these quantities have not been directly measured, the predicted decrease in time constants point to networks becoming more responsive over time, while reduced input may reflect the disappearance of spontaneously active subplate neurons later in development [77–79]. We also note that, in contrast to measurements of increased membrane capacitance in organoids [80], model capacitance on average decreases over time (Supplemental Fig. S4a), highlighting that model parameters should be interpreted as *hypotheses* for mechanistic explanations.

Next, while the ensemble of model configurations tightly reproduce burst features in each target recording, many parameters appear to have a large range of possible values. However, degenerate parameters exhibit relationships that reveal much tighter constraints: For example, across 100 discovered models of an earlier recording (week 9, blue), there is a weak but significant relationship between adaptation current strength and time constant (Fig. 3e). At a later timepoint (week 32, orange), not only do both parameters shift drastically in values, their correlation also becomes much stronger, exposing a narrow subspace on which fast network bursting behavior is invariant.

Similarly, the ratio of total excitatory to inhibitory synaptic charge (E:I charge) has a strong negative correlation with the E:I ratio of synaptic count in earlier recordings (see Table 3 for definitions). That is, early networks typically have weak inhibition, and either stronger or greater numbers of excitatory synapses, but not both, in order to preserve the same network bursting dynamics (Fig. 3f). At the later time point, however, networks on average shift to a more inhibition-balanced regime while decoupling from synaptic count ratio. With stronger inhibitory synaptic charges (leftwards on x-axis), there is a lesser reliance on balancing excitatory connection strength and count. Thus, by leveraging efficient model discovery, we uncover a variety of trajectories in correlations between model parameters at each timepoint (Fig. 3g)— even observing sign changes—suggesting strong and dynamic-dependent invariances over the course of development (see Supplemental Fig. S4c for other parameters).

### *In silico* perturbation reflects *in vitro* experiments and refines candidate mechanisms

Given the ensembles of dynamic-consistent models, we can further narrow down candidate circuit mechanisms via perturbation experiments. We demonstrate this through a pharmacological manipulation: Bicuculline, a GABA_A_ antagonist, was applied acutely after a baseline recording at week 27, which increased overall activity and decreased inter-burst intervals (Fig. 4a). We emulate this perturbation *in silico* by setting inhibitory synaptic strengths to zero (*Q*_*I*→*E*_ and *Q*_*I*→*I*_) in 100 discovered models that matched network activity of the baseline recording.

**Figure 4.**
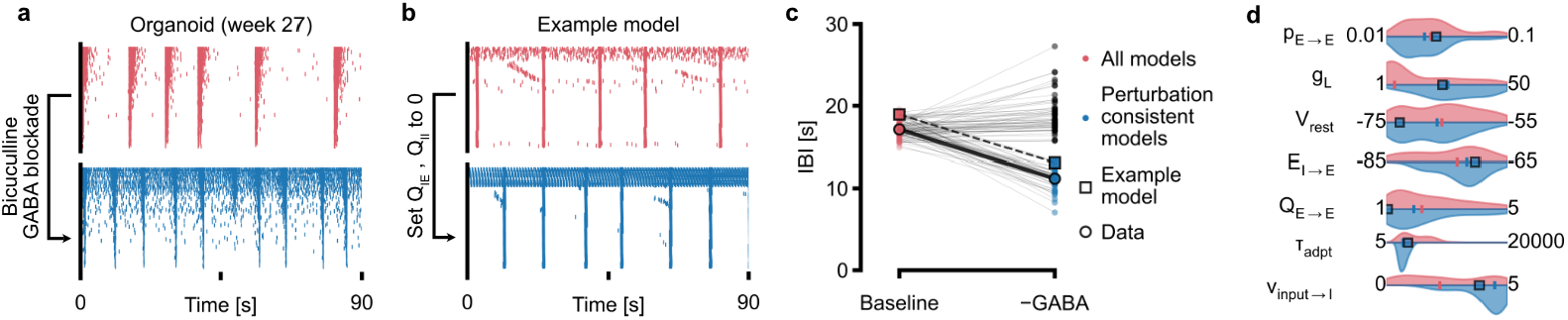
Causal perturbation of models *in silico* reproduces GABA_A_ blockade in organoids. **a)** Organoid recording at week 27, before (top) and after (bottom) application of 5 mM/mL bicucculine (GABA_A_ antagonist). **b)** Example discovered model of the baseline recording (top), and the same model after inhibitory synaptic strengths are set to 0 (*Q*_*I→E*_ and *Q*_*I→I*_). **c)** A subset of the 100 dynamic-consistent baseline models further reproduce the effect of GABA blockade in organoids (decreased IBI) after *in silico* perturbation. **d)** Example parameter distributions of all baseline models (pink) vs. only those that are also consistent with the perturbation recording (blue). See Supplemental Fig. S5 for all parameters.

Re-simulating the now inhibition-depleted models, a subset of them produced the similar behavior of reduced IBI (example model in Fig. 4b), while others exhibited unchanged or even increased IBI (Fig. 4c). What differentiates between models that are consistent from those that are inconsistent with dynamics during GABAergic blockade? Examining the parameters of only the perturbation-consistent models, we see that the distribution of many parameters were essentially unchanged (Fig. 4d, Supplemental Fig. S5). However, a few were more constrained, such as adaptation time constant (*τ*_*adpt*_) and inhibitory background input (*ν*_*input*→*I*_), suggesting that their values are more important for realistic models of organoid network dynamics. Thus, we demonstrate how *in silico* perturbation experiments can further refine candidate models and mechanistic hypotheses.

### Inverse modeling of mouse cortical and hippocampal circuit dynamics

Next, we showcase the flexibility of AutoMIND and discover circuit models of Neuropixels recordings from mouse hippocampus and visual cortex (Fig. 5a, data from [52]). During spontaneous behavior, individual neurons exhibit sparse and irregular firing across subregions of the visual cortex and hippocampus (Fig. 5b). We compute the population firing rate for each subregion, which appears largely asynchronous and noise-like (Fig. 5c). However, a variety of commonly observed dynamics are revealed in the frequency domain, such as high-and low-frequency, narrowband oscillations, as well as broadband 1/f-like patterns (Fig. 5d, black). In contrast to approaches that only target narrowband features such as gamma power, here we aim to quantitatively reproduce the *entire* power spectral density (PSD) up to 495 Hz (i.e., 990 spectral power features).

**Figure 5.**
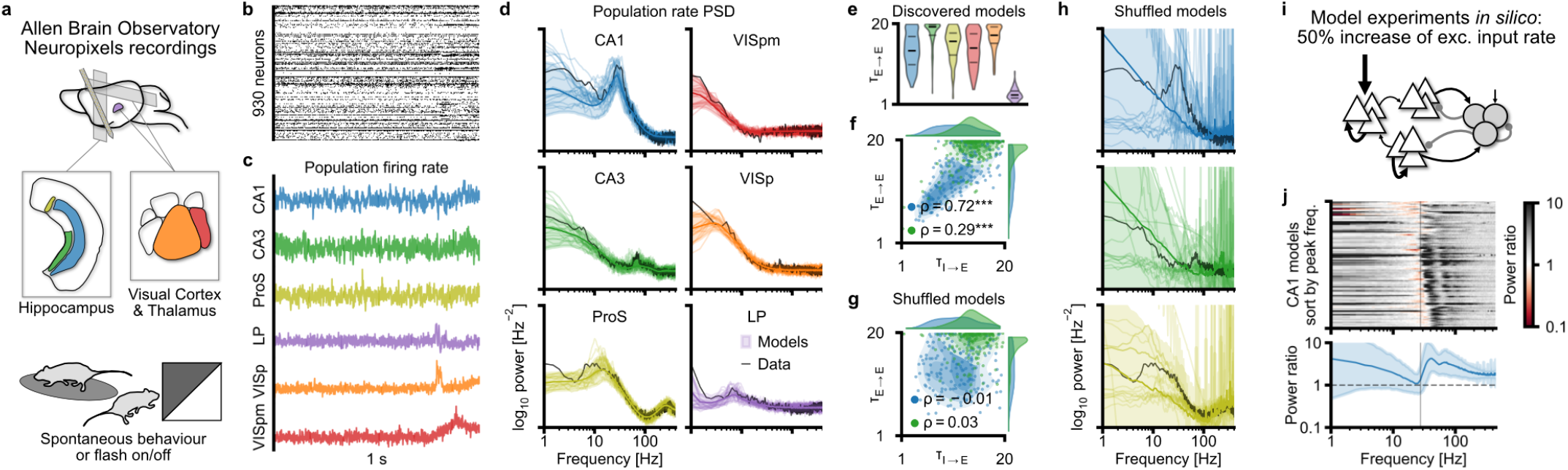
Hippocampal and cortical circuit models capture diverse spectral profiles of region-specific population dynamics. **a)** Neuropixels recordings across mouse visual and hippocamal areas from the Allen Brain Observatory Visual Coding dataset [52]; analyses restricted to periods of spontaneous behavior or black/white screen trials. **b)** 1-second example spike raster of simultaneously recorded neural population. **c)** Population firing rate of cells collapsed across subregions. ProS: prosubiculum, VISpm/VISp: posteriomedial/posterior visual cortex, LP: lateral posterior nucleus (of the thalamus) **d)** Population rate power spectral density (PSD) of real recordings (black) and simulations of discovered models (colored); thin lines show 20 example models, thick lines and shaded regions denote mean *±* standard deviation across 200 data-consistent models. **e)** Example parameter distributions across regions, lines denote median and quartiles. **f)** Parameter correlations across CA1 vs. CA3 models. Spearman correlation, *** denotes p<0.001. *tau*_*E→E/I→E*_ : decay time constant of E-E / I-E synapses. **g)** Independently shuffling parameter values of models in each region destroys pairwise correlation (scatter plot) while retaining individual parameter distributions (histograms, top and right). **h)** Simulations of shuffled models vary drastically and fail to reproduce target network dynamics. **i)** Schematic of *in silico* experiment: 50% increase in external input rate to excitatory cells. **j)** (Top) Spectral response (post:baseline power ratio) of individual CA1 models under increased input; models sorted by peak frequency of baseline model, vertical line denotes peak frequency of data (30 Hz). (Bottom) Average power change across all CA1 models shows increase in peak frequency and power, and broadband power; mean *±* standard deviation of *log*_10_power ratio. See also Supplemental Fig. S6-S9.

Importantly, due to the flexibility of the clustered-AdEx network, we can leverage the *same* mechanistic model—and thus reuse the *same* training simulations—to study diverse frequency profiles of population activity across brain regions. To do so, we similarly compute from network simulations the population rates and their normalized PSDs (e.g., Fig. 2d). Then, we train a new DGM to map parameters to simulated network activity, which now resides in a much bigger, 990-dimensional observation space. Finally, we query the DGM for candidate models, whose simulated dynamics have spectral profiles that closely match the observed PSDs (Fig. 5d; Supplemental Fig. S6 for all observations).

In 200 dynamic-consistent models of each region, we again find that many parameters take on a broad range of possible values (Fig. 5e; Supplemental Fig. S7a for all parameters). Similarly, there are significant correlations between parameters, such as the time constants of excitatory and inhibitory synapses (Fig. 5f; Supplemental Fig.. S7b). Moreover, specific pairwise correlations across invariant models differ for different target regions (e.g., CA1 vs. CA3), demonstrating again the dependence of parameter invariances on dynamics.

These findings indicate that while exact values for individual parameters can vary broadly, the geometry of invariance defined by their pairwise and higher-order relationships may be critical. To test this, we independently shuffled the values of each parameter within models for each region, such that individual parameter distributions match those of the original models but higher-order relationships are destroyed (Fig. 5g). Interestingly, the majority of shuffled models still produce valid simulations, but very few reproduce the original target frequency profiles (Fig. 5h; Supplemental Fig. S8). Thus, individual parameter ranges may define a broad “viable” zone of realistic network activity, but the precise target dynamics is only captured by models within the invariant parameter subspace.

Finally, while discovered models are degenerate in their parameter configurations, we observe a surprising functional equivalence and robustness in their response to perturbation. We set up an *in silico* experiment where the excitatory input rate of discovered models is increased by 50% from their original value (Fig. 5i). Based on existing studies on gamma oscillatory dynamics in network models [81, 82], we hypothesized that increased excitatory drive would lead to an increase in oscillatory frequency and power, though it is unclear whether all models would respond as such.

Surprisingly, almost all CA1 models showed reduced power at their original oscillatory peak frequency of 30 Hz and increased power at around 35-45 Hz, in addition to an overall increase in broadband power, which are visible when averaged across models as well (Fig. 5j). Furthermore, when a clear oscillation was present, models of visual and hippocampal areas almost always exhibited a shift towards higher frequency—regardless of the baseline frequency— as well as a broadband power increase that was especially prominent in high frequencies (Supplemental Fig. S9). Interestingly, models differed in their low-frequency response and in an area-dependent manner, where some areas instead showed a decrease in power consistent with network de-synchronization (e.g., ProS vs. VISp, Supplemental Fig. S9). Thus, we show how AutoMIND enables the discovery of invariant parameter configurations underlying complex frequency profiles of population dynamics across the mouse brain, and accelerates hypothesis testing *in silico*.

### Dynamic-dependent global invariances and local sensitivity in circuit models

From both experimental datasets, we discovered dynamic-consistent models with degenerate parameters, extending previous studies on single-neuron and small circuit dynamics [20, 22]. Furthermore, using a common spiking network to model vastly different dynamics revealed that parameter invariances in degenerate configurations are dynamic-dependent. Are such degeneracies only a reflection of uncertainties due to potential mismatches between the model and the underlying circuit, or are dynamic-dependent invariances a general property of neural circuit models?

To systematically address this, we chose a diverse set of held-out simulations as synthetic target observations from “*in silico* circuits” (Fig. 6a, Supplemental Fig. S2). Thus, we guarantee that target observations can be perfectly reproduced by the model, while having access to the “true” parameters used to generate them. We again target the entire power spectrum of the population firing rate, where we can directly use the previous DGM without retraining.

**Figure 6.**
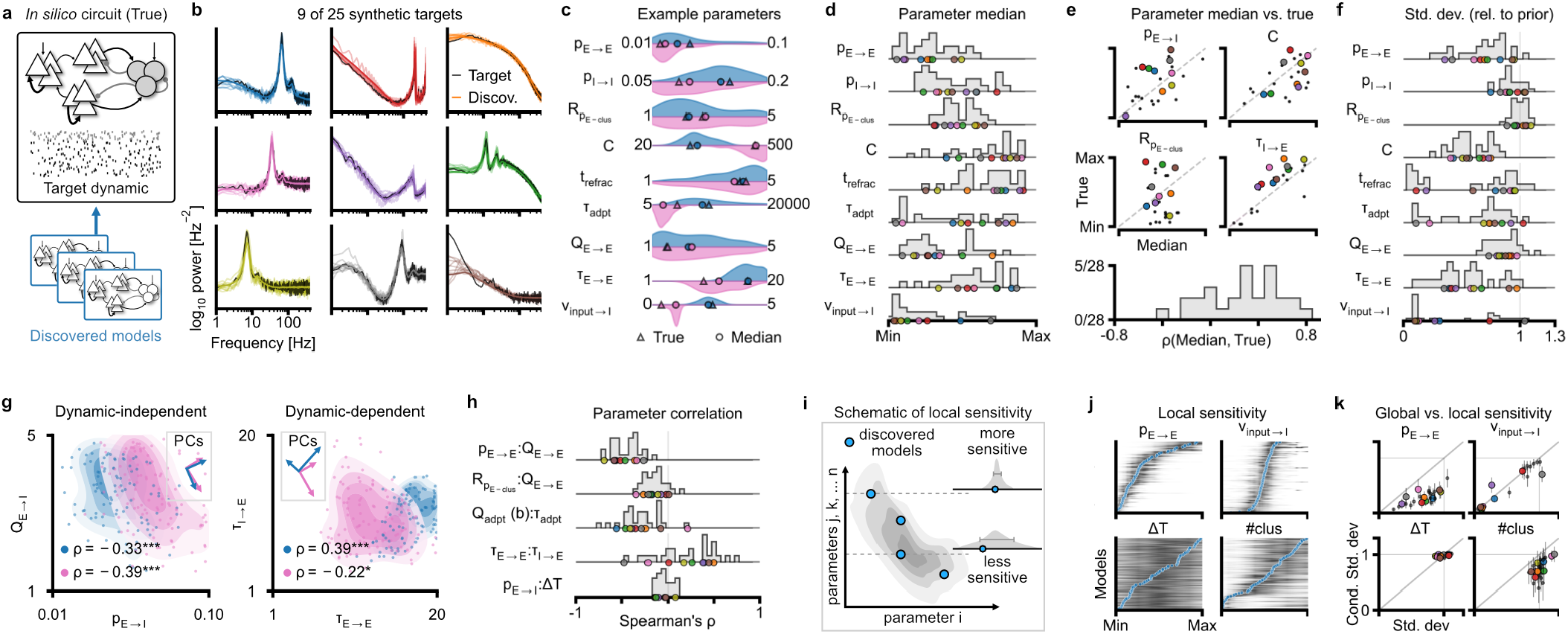
Dissecting global parameter invariance and local sensitivity across dynamical regimes. **a)** Schematic: discovering models (colored) consistent with synthetic target observations from “*in silico* circuits” (black) with known simulation parameters. **b)** Population firing rate PSDs of 9 example observations (black), and of the discovered models (colored; thin lines show 10 individual models, thick line and shaded region denote mean *±* std. across 100 models). Same colors used in **(c-h). c)** Example parameter distributions for two target observations exhibiting gamma-range oscillation. Violins over 100 models, circles denote medians of distribution, triangles denote parameter values used to generate the observation (True). **d)** Parameter medians of discovered models; x-axis re-scaled to parameter range, histograms taken over medians from each of the 25 targets, colored circles correspond to example targets in **(b). e)** Correlation between true and median value of discovered models over 25 targets, for 4 parameters (top), and histogram over all parameters (bottom); Spearman correlation. **f)** Parameter standard deviations of discovered models; x-axis re-scaled such that the standard deviation of the uniform prior distribution is 1. **g)** Consistently negative correlation between synaptic connection probability and strength (left), and inconsistent correlation between excitatory and inhibitory synaptic time constant (right), for two different target dynamics. Spearman correlation, */**/*** denote p=0.05, 0.01, and 0.001 respectively. Inset: schematic of aligned (left) vs. unaligned (right) Principal Components between blue and pink datapoints. **h)** Example parameter correlations in discovered models for all 25 targets (histogram), showing dynamic-independent and dynamic-dependent correlations. **i)** Schematic of sensitivity analysis: at each discovered model, the value of one parameter is re-sampled while holding the others constant; repeat at all locations (models). **j)** Example parameters with different local sensitivity (black shading denotes conditional probability density); *p*_*E→E*_ can take on a broad range of values across discovered models (blue dots), but is *locally sensitive* given the values at each discovered model; *ν*_*input→I*_ is both globally and locally sensitive, while Δ*T* is insensitive; #*clus* is sensitive at low values (one or two clusters), but insensitive at higher number of clusters. **k)** Parameter standard deviations of discovered models (same as **(f)**) vs. standard deviation of conditional probability densities from **(j)** (mean *±* std) for four example parameters and all target observations. Diagonal line denotes identity. See Supplemental Fig. S10–S13 for all observations and parameters.

First, AutoMIND discovers ensembles of models that closely match synthetic target observations (Fig. 6b, Supplemental Fig. S10). Even for relatively similar dynamics, like different narrowband, high-frequency oscillations (Fig. 6b, first column), model parameters exhibit a range of behaviors (Fig. 6c) from broadly distributed (e.g., *Q*_*E*→*E*_) to tightly constrained (e.g., *ν*_*input*→*I*_). Across 25 target observations, median parameter values of ensembles span nearly the full range (Fig. 6d, Supplemental Fig. S11a), indicating that a large volume of parameter space is required to capture different dynamical regimes. However, discovered parameter configurations do not in general match the “true” values used in generating the target observation (Fig. 6e, Supplemental Fig. S11c). Furthermore, the variability of parameters differs greatly across target dynamics (Fig. 6f, Supplemental Fig. S11b): Some parameters are no more constrained than the prior uniform distribution (e.g., 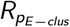), while others strongly depend on the target dynamic, and can in some cases be almost perfectly identified (e.g., *τ*_*adapt*_, *ν*_*input*→*I*_). Thus, even for target observations generated by the same model with known parameter values, dynamic-consistent configurations exhibit strong degeneracy.

Can we similarly detect invariance structures across degenerate models? Again, we consistently observe strong correlations between parameters. In some cases, the invariance is intuitive: the probability of synaptic connection (*p*_*E*→*I*_) scales inversely with connection strength (*Q*_*E*→*I*_) to preserve the same network dynamic (Fig. 6g, left), and this is likely to be consistent regardless of the target dynamics. However, we also see dynamic-dependent correlations, such as between excitatory vs. inhibitory synaptic time constants (Fig. 6g, right). Overall, we find a variety of behaviors (Fig. 6h, Supplemental Fig. S11c) from consistently strong and negative correlations (*p*_*E*→*E*_ :*Q*_*E*→*E*_), consistently weak but significant 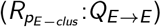, to no correlation at all (*p*_*E*→*I*_ :Δ*T*), in addition to strong dependencies on target dynamics.

Geometrically, the dependence of parameter correlations on the target observation implies that invariant models for different dynamics reside on different lower-dimensional (nonlinear) subspaces (insets of Fig. 6g). If so, the space encapsulating all dynamic-consistent models—being the union—should necessarily be higher dimensional than that for each individual one. We confirm this with Principal Component Analysis: fewer PCs are required to explain the same proportion of variance when fit to the ensemble of models for the same target observation, than PCs fit to the same number of models randomly sub-sampled across ensembles (Supplemental Fig. S12a). We further confirm the necessity of these higher-order relationships by independently shuffling the values of each parameter within the same ensemble, where shuffled models no longer reproduce the target dynamics (Supplemental Fig. S12b). Thus, there exists for each target observation a *different* subspace on which network dynamics is invariant, typically spanning a moderate volume of the entire parameter space (when measured by linear dimensionality).

Finally, we showcase how AutoMIND enables characterization of the local sensitivity (or stiffness [25]) along invariant subspaces. This corresponds to the question: “holding all other parameters j,k,l… constant, how much can parameter i vary?” To answer this, we first define a “location” at a single parameter configuration, holding the values of all parameters fixed except one, and re-sample that parameter from the 1-dimensional conditional distribution, *p*(*θ*_*i*_ |*x, θ*_*j*_, *θ*_*k*_ …). If this parameter is locally sensitive, then only a narrow range of values is possible, thus producing a thin distribution (i.e., low standard deviation, schematic in Fig. 6i). Conversely, a broader distribution with high standard deviation implies that the target dynamics is less sensitive to perturbations along this direction, at this location.

Looking first only at a single target observation (top left in 6b), we see that while a parameter, such as *p*_*E*→*E*_, can take on a broad range of values across discovered models (blue points in Fig. 6j), it is tightly constrained given the values of the other 27 parameters (i.e., locally sensitive). In contrast, some parameters (e.g., Δ*T*) always have a full range of variation, even when holding all others constant, which means that they are largely inconsequential. Lastly, for some parameters like the number of excitatory clusters (#*clus*), we observe *location-dependent sensitivity* : discovered models can vary broadly in the number of clusters, but models with fewer clusters (one or two) are more tightly constrained to that value, while models with more clusters have more “slack”. Across all target observations, we see that the extent of a parameter’s location-dependent sensitivity, relative to its overall range of variation, also differs for different dynamics (Fig. 6k, Supplemental Fig. S13). Thus, AutoMIND reveals dynamic-dependent global parameter invariances, as well as dynamic-and location-dependent sensitivity of the subspace, shedding light on how circuit mechanisms cooperate to maintain stable dynamics in spiking neural network models.

## Discussion

We present AutoMIND, a deep inverse modeling framework for efficient discovery of mechanistic model configurations that are quantitatively consistent with target observations of neural dynamics. This is jointly enabled by an expressive spiking network model that exhibits a rich variety of experimentally observed dynamics, in combination with probabilistic deep generative models and simulation-based inference. We demonstrate on a number of experimental and synthetic datasets that many parameter configurations are consistent with a single observation, and that ensembles of candidate models are valuable for interpreting circuit mechanisms and testing causal perturbations *in silico*. Moreover, models exhibit parameter invariances with global and local structures that are, in contrast to intuitive expectation, dependent on the target dynamics. Our method and findings have broader utility and implications for experimental and computational investigations of neural circuit mechanisms underlying population dynamics.

### Direct and extended application of AutoMIND

The inverse modeling framework is flexible and, together with the trained deep generative models, model simulations, and code repository we make publicly available, can be customized for a variety of applications:

First, to discover configurations of the same network model while targeting the summary features presented here, we provide trained DGMs that can be directly queried for new data, with no further network optimization or simulations required. This is enabled by the amortization property of neural posterior estimation [24, 49, 83] and requires minimal additional programming, while discovered configurations can be simulated to further refine candidate models based on consistency with experimental data. Second, to discover configurations of the same spiking network but targeting other summary features, it is only necessary to compute them from the provided database for training new deep generative models. Our method can be extended to account for other recording modalities as well, including local field potentials (LFP), magneto/electroencephalography (M/EEG), optical imaging, and even functional magnetic resonance imaging (fMRI), as long as a forward model of how the signal is generated from population spiking is provided [84, 85].

In both cases, AutoMIND can drastically accelerate the process of discovering dynamic-consistent models while facilitating direct parameter comparisons for the *same* spiking network, and therefore the encoded mechanisms across different studies. Another important use case of the existing database of simulations—both the training set and discovered models—is parameter space analysis and perturbation testing, where *hundreds* of models that quantitatively reproduce the same dynamic, such as network oscillations of a specific frequency or scale-free behavior, are readily available.

Finally, to investigate additional circuit mechanisms, either by making fixed parameters free in the current model (e.g., inhibitory cell parameters) or by introducing new mechanistic models altogether, a new database of training simulations and the associated summary features are necessary for training new DGMs, but no experimental data is required at any point in this process. More generally, the current study, along with several recent others [24, 44–48], highlight the utility of training (deep) generative models on model simulations to invert the data generation mechanisms underlying real neural recordings at different resolutions.

### Limitations of the mechanistic and generative models

“*All models are wrong, but some are useful*” [86]. While the clustered-AdEx network model proposed here exhibits a wide range of dynamics, it is neither the “correct”, nor a universal model of circuit dynamics. Indeed, numerous mechanisms are not encoded as either free or fixed parameters in the current network: For example, it lacks diverse inhibitory classes, which have been shown to be critical for recapitulating aspects of e.g., gamma oscillations *in vivo* [87], and could be represented by additional inhibitory populations [44, 88]. Further examples include dendritic morphology, active and passive conductances with different timescales (e.g., calcium), laminar structures, layer-dependent and structured cellular and connectivity profiles, and short-term and long-term plasticity mechanisms, all of which likely play an important role in neural population dynamics and computation, but are inconsistently accounted for across existing models of neural circuits.

As a consequence, while the model can faithfully reproduce temporal dynamics of neural populations in experimental recordings, it can also exhibit unrealistic behavior in single-neuron firing statistics. This may in part be caused by a lack of heterogeneity in the model beyond clustered connectivity, which was critical for introducing variability in spatial and temporal dynamics. These observations point to a mismatch between the model and real neural circuits. Such model misspecification may be ameliorated by potential extensions to the spiking network [89, 90], as well as misspecification-aware inference algorithms [91–93], which can be easily incorporated into the inverse modeling framework and are likely to induce greater parameter degeneracy.

On the other hand, even given the misspecified model, it is possible that the current parameter space has not been fully explored, and that other dynamic-consistent configurations may exist in entirely different regions. This can be mitigated with a larger training simulation dataset, as well as by incorporating more powerful deep generative models, such as diffusion models [94]. With greater coverage of the parameter space, it is possible that the precise structure of parameter invariances will change, but likely not their observed dependence on dynamics.

### Implications of dynamic-dependent invariances in circuit mechanisms

The observation of dynamic-dependent invariance raises a number of questions with fundamental implications for studying circuit mechanisms underlying neural population dynamics:

First, in the subspace of model configuration for a particular dynamical regime, where does the real circuit sit? Reproducing the population dynamics of a specific target observation is just one constraint, while biological constraints faced by neural circuits *in vivo* may further shrink the viable subspace of possible models. For one, the same circuit can typically exhibit distinct dynamical regimes at different times, such as asynchronous and synchronous activity. Does this imply that neural circuits must exist at the intersection of the corresponding invariant subspaces, or is it sufficient to be “in the neighborhood”, such that hopping from one to the other is a possible means of switching between dynamical states? In either case, it is unclear to what extent such a change is driven solely by network inputs, versus factors that change circuit properties on a longer timescale, such as diffuse sub-cortical neuromodulators [28, 95]. Furthermore, degeneracy and sloppiness have long been argued to be not just a byproduct of stochasticity, but meaningful variability that provides robustness for functionality and survival [17–19, 21, 96]. In this vein, dynamic flexibility and stability may be additional constraint, especially when computationally relevant [97, 98], in addition to fitness constraints such as metabolic cost [30]. Such multi-objective optimization may result in preferable model configurations on the “Pareto front” [31, 99].

Second, and on a more cautionary note, the consistency with which we observe model degeneracy suggests that care should be taken when making mechanistic conclusions from *individual* (or a single pair of) dynamic-consistent model configurations. Parameter distributions of discovered models often have long, overlapping tails. In these cases, it is possible to detect a difference from a pair of models opposite to that of the ensembles on average (e.g., Fig. 6c *τ*_*E*→*E*_). Without ruling out the existence of such statistically unlikely explanations, it is important to interpret them with caution, in addition to performing perturbation experiments *in vivo* or *in silico* to further isolate causal mechanisms. This highlights the importance of ensemble modeling, and efficient methods for doing so.

Finally, parameter invariances and higher-order correlations call into question the conceptual utility of “single-factor mechanisms” for understanding neural circuit dynamics, as well as for modeling neurological disorders [31]. While it is possible to perform single-factor perturbation experiments *in silico* and *in vivo*, and to observe such isolated causal mechanisms from natural variation (e.g., single-nucleotide polymorphisms), the ubiquity of degeneracy argues against such simplistic conceptualization, especially in robust biological systems. Our results suggest that “circuit mechanisms” may be better conceptualized as invariances and the changes thereof, similar to the mechanistic manifold interpretation of neural dynamics [13, 97], rather than a difference in individual parameters between state A and B. Moreover, invariances reveal opportunities to coarse-grain microscopic physical variables into fewer phenomenological variables while maintaining the same explanatory power [33, 100]. However, the dynamic-dependence of invariances poses further challenges for designing general-purpose models of neural dynamics at the appropriate resolution, since how microscopic variables can be reduced would depend on the specific dynamic observed, suggesting the necessity of state-dependent “meta” models.

## Acknowledgments

RG was supported by the European Union’s Horizon 2020 research and innovation program under the Marie Skłodowska-Curie grant agreement No. 101030918 (AutoMIND). This work was supported by the German Research Foundation (DFG) through Germany’s Excellence Strategy (EXC-Number 2064/1, PN 390727645) and SFB1233 (PN 276693517), and the German Federal Ministry of Education and Research (Tübingen AI Center, FKZ: 01IS18039), and the European Union (ERC, “DeepCoMechTome”, ref. 101089288). MD and AS are members of the International Max Planck Research School for Intelligent Systems (IMPRS-IS). The authors thank Mattia Chini, Randolph Helfrich, Oleg Vinogradov, Anna Levina, Alysson Muotri, Guy Moss, and members of Mackelab for discussions throughout the project.

## Methods

### Code and data availability

All code, including for generating results and figures of this publication, as well as data repository links for the trained deep generative models, raw, and processed simulation data, are available at https://github.com/mackelab/automind.

### Clustered adaptive exponential integrate-and-fire spiking neural network model

In this section, we detail the equations governing the dynamics of the adaptive exponential integrate-and-fire (AdExIF) neuron model and for the conductance-based synapse. Then, we describe how networks are connected, including the cluster formation procedure, as well as external input connections. We summarize the free parameters of the model and their possible ranges, in addition to the values of important fixed parameters. In the following equations, variables in blue denote free parameters on which inference is performed.

#### Membrane potential and adaptation current dynamics

Membrane voltage (*V*) and adaptation current (*w*) dynamics are governed by the AdExIF differential equations from Brette and Gerstner [53]:

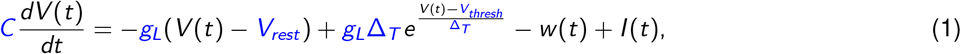

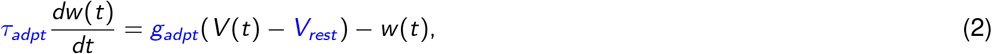

where:

- *V* (*t*) and *w* (*t*) are state variables representing the membrane voltage and adaptation current, respectively
- *I* (*t*) is the total input current from recurrent and external inputs
- *C* is the membrane capacitance
- *g*_*L*_ is the leak conductance
- *V*_*rest*_ is the resting membrane potential
- Δ_*T*_ is the slope factor
- *V*_*thresh*_ is the threshold potential
- *τ*_*adpt*_ is the time constant of the adaptation current
- *g*_*adpt*_ is the sub-threshold adaptation coupling parameter.

Action potential firing is instantaneous, i.e., when *V* (*t*) reaches threshold voltage at time *t*_*f*_, *V* (*t*) ≥ *V*_*thresh*_:

- The spike time *t*_*f*_ is recorded
- The membrane potential *V* (*t*) is reset: *V* (*t*_*f*_) ← *V*_*reset*_
- The adaptation current *w* (*t*) is increased by a fixed amount: *w* (*t*) ← *w* (*t*_*f*_) + *Q*_*adpt*_

Note that *g*_*adpt*_ and *Q*_*adpt*_ refer to the variables typically labeled as *a* and *b* [53], but are renamed here for clarity, as they represent a conductance-like sub-threshold coupling term and a spike-triggered quantal increase of adaptation current, respectively.

Both excitatory and inhibitory neurons follow these dynamics, but the variables marked in blue above are free parameters for excitatory neurons only (Table 1), while most parameter values for inhibitory neurons are fixed (Table 2).

**Table 2.** Fixed parameters of the clustered AdEx network model (many adapted from Zerlaut et al. [63]).

| Parameter | Explanation | Value | Unit |
| --- | --- | --- | --- |
| *Network parameters* |  |  |  |
| $\%N_{total}$ | Number of neurons | 2000 | unitless |
| *Cellular parameters for inhibitory neurons* |  |  |  |
| $C$ | Membrane capacitance | 200 | $pF$ |
| $g_L$ | Leak conductance | 10 | $nS$ |
| $V_{rest}$ | Resting potential | -65 | $mV$ |
| $V_{thresh}$ | Threshold potential | -50 | $mV$ |
| $V_{reset}$ | Reset potential | -65 | $mV$ |
| $t_{refrac}$ | Refractory period | 5 | $ms$ |
| $\Delta_T$ | AP slope factor | 2 | $mV$ |
| $g_{adpt}(a)$ | Subthreshold adaptation coupling | 0 | $nS$ |
| $Q_{adpt}(b)$ | Spike-triggered adaptation quantal current (b) | 0 | $pA$ |
| $\tau_{adpt}$ | Adaptation time constant | 500 | $ms$ |
| *Synapses onto excitatory neurons* |  |  |  |
| $E_{E \rightarrow E}$ | Excitatory to excitatory synapse reversal potential | 0 | $mV$ |
| *Synapses onto inhibitory neurons* |  |  |  |
| $E_{E \rightarrow I}$ | Excitatory to inhibitory synapse reversal potential | 0 | $mV$ |
| $E_{I \rightarrow I}$ | Inhibitory to inhibitory synapse reversal potential | -80 | $mV$ |
| $\tau_{E \rightarrow I}$ | Excitatory to inhibitory time constant | 5 | $ms$ |
| $\tau_{I \rightarrow I}$ | Inhibitory to inhibitory time constant | 5 | $ms$ |
| *External input* |  |  |  |
| $N_{Poisson \rightarrow E/I}$ | Number of Poisson synapses (onto E or I neurons) | 500 | unitless |
| $Q_{Poisson \rightarrow E/I}$ | Poisson synapse release conductance (onto E or I neurons) | 1 | $nS$ |

At the start of simulation, membrane voltages for all neurons are randomly initialized according to a Gaussian distribution, *V* (0) ∼ N(*V* ; *µ*=*V*_*rest*_, *σ*=4), i.e., centered at *V*_*rest*_ with standard deviation 4, while *w* (0) = 0.

#### Synapse dynamics

Excitatory and inhibitory synapses are conductance-based and follow the (“single-exponential”) differential equation,

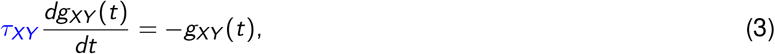

and the current as a result of the conductance at a single synapse is

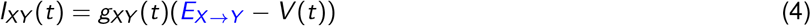

where:

- *X* and *Y* can be *E* or *I*, for excitatory or inhibitory, respectively: *X, Y* ∈ {*E, I* }
- *g*_*XY*_ (*t*) and *I*_*XY*_ (*t*) are the state variable representing conductance and current from presynaptic population X onto a cell in postsynaptic population Y
- At presynaptic spike time *t*_*f*_, the postsynaptic conductance is increased by a fixed amount, with a possible delay:
- *g*_*XY*_ (*t*) ← *g*_*XY*_ (*t*_*f*_ + *τ*_*delay*_) + *Q*_*X*→*Y*_ (though we keep *τ*_*delay*_ = 0 in all simulations)
- *τ*_*X*→*Y*_ is the time constant of the conductance
- *E*_*X*→*Y*_ is the reversal potential of the synapse
- The total synaptic current onto a single neuron is the summation over all its incoming synapses, i.e., Σ*I*_*XY*_ (*t*).

#### Clustered connectivity for heterogeneous networks

Networks always consist of *N*_*total*_ = *N*_*exc*_ + *N*_*inh*_ = 2000 neurons in total, but with the proportion of excitatory neurons %*E* :*I* a free variable, i.e., *N*_*exc*_ = *N*_*total*_ (%*E* :*I*) excitatory neurons and *N*_*inh*_ = *N*_*total*_ (1 − %*E* :*I*) inhibitory neurons. Binary connections between neurons are drawn from a Bernoulli distribution with probability *p*_*X*→*Y*_, where *X, Y* ∈ {*E, I* }. In other words, the connection matrix *W*_*XY*_ with elements *w*_*ij*_ ∼ Bernoulli(*p*_*X*→*Y*_) is a random binary matrix, and is sampled within `Brian2` for a given random seed. If a connection exists, a synapse is instantiated with an effective weight of *Q*_*X*→*Y*_.

The connectivity described thus far would form four homogeneous (Erdő s–Rényi) sub-graphs between *E* →*E, E* →*I, I* →*E*, and *I* →*I* populations depending on the respective *p*_*X*→*Y*_. However, *E* →*E* connectivity is clustered and formed via a procedure similar to Litwin-Kumar and Doiron [57]: First, the number of clusters (#*clus*, a free parameter) is specified. Then, each excitatory neuron is assigned to 2 of the #*clus* clusters. Neurons that share cluster membership have increased connection probability 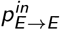 and strength 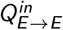 determined by two amplification parameters, where 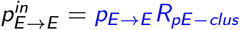 and 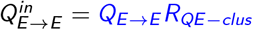. Connections are randomly drawn as above, but with 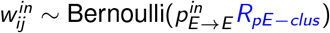. Connections between excitatory neurons that are not in the same cluster, and all connections to and from inhibitory neurons, are homogeneous and follow the respective non-amplified probabilities.

#### External Poisson input

Neurons receive external Poisson input that mimic spontaneously occurring activity or upstream populations not represented in the model. External input onto an individual neuron is instantiated as *N*_*poisson*_ = 500 independent Poisson synapses (identical to the conductance-based synapse described above) with a fixed quantal conductance *Q*_*poisson*_ = 1*nS*. Each Poisson synapse has as firing rate *ν*_*input*→*E*_ or *ν*_*input*→*I*_ depending on its downstream population (E or I), which are free parameters shared between all neurons, while spikes are drawn from independent homogeneous Poisson processes regardless of whether they share a downstream target. Finally, to further introduce heterogeneity in network activity, only a subset of randomly chosen excitatory neurons—determined by the free parameter %*input*_*E*_ — receive input.

#### Free and fixed parameters

In total, the network model has 28 free parameters whose values are either randomly sampled (in the training dataset) or inferred given a target observation. These parameters are summarized in Table 1, including their bounds (within which parameters are sampled uniformly), and physical units. A number of other important model parameters are fixed, summarized in Table 2. Finally, “derived” parameters, such as those presented in Fig. 3 (e.g., *τ*_*RC*_), are computed as a function of the free parameters, and their definitions can be found in Table 3. Note that when computing synaptic excitation and inhibition properties across the network, we only consider synapses onto excitatory neurons, reflecting the measurement of synaptic input balance in pyramidal neurons.

**Table 3.**
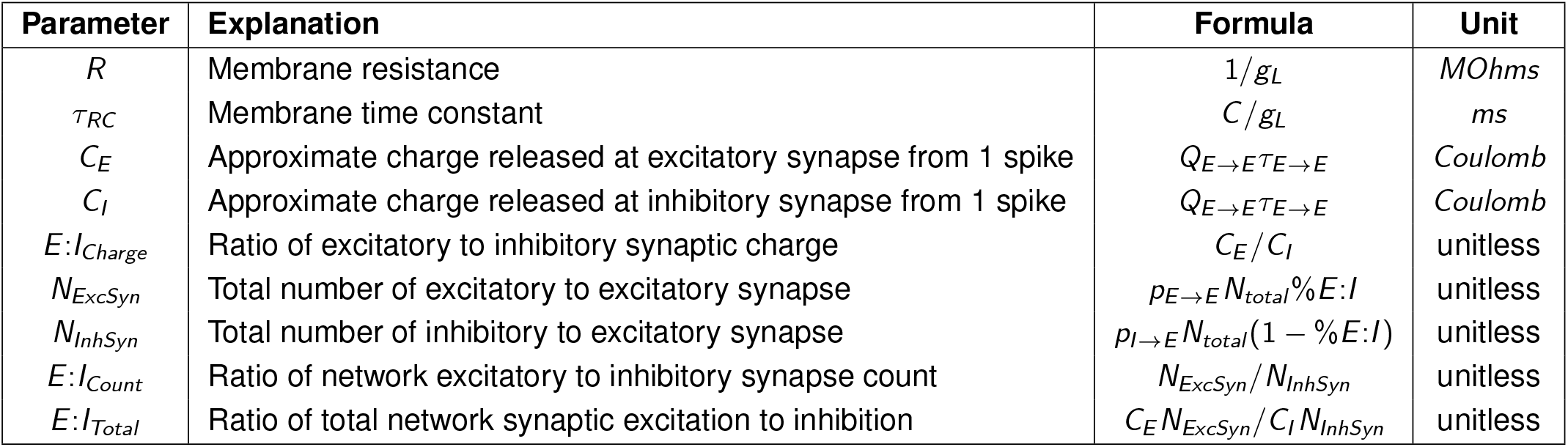
Derived parameters computed as a function of the free parameters in Table 1.

| Parameter | Explanation | Formula | Unit |
| --- | --- | --- | --- |
| $R$ | Membrane resistance | $1/g_L$ | *MOhms* |
| $\tau_{RC}$ | Membrane time constant | $C/g_L$ | *ms* |
| $C_E$ | Approximate charge released at excitatory synapse from 1 spike | $Q_{E \rightarrow E} \tau_{E \rightarrow E}$ | *Coulomb* |
| $C_I$ | Approximate charge released at inhibitory synapse from 1 spike | $Q_{E \rightarrow E} \tau_{E \rightarrow E}$ | *Coulomb* |
| $E:I_{Charge}$ | Ratio of excitatory to inhibitory synaptic charge | $C_E/C_I$ | unitless |
| $N_{ExcSyn}$ | Total number of excitatory to excitatory synapse | $p_{E \rightarrow E} N_{total} \% E:I$ | unitless |
| $N_{InhSyn}$ | Total number of inhibitory to excitatory synapse | $p_{I \rightarrow E} N_{total} (1 - \% E:I)$ | unitless |
| $E:I_{Count}$ | Ratio of network excitatory to inhibitory synapse count | $N_{ExcSyn}/N_{InhSyn}$ | unitless |
| $E:I_{Total}$ | Ratio of total network synaptic excitation to inhibition | $C_E N_{ExcSyn}/C_I N_{InhSyn}$ | unitless |

### Network simulations

Here, we describe model simulation settings, early stopping for the simulations, and simulation quality assurance criteria, as well as simulation data storage and preprocessing steps.

#### Simulation settings and early stopping

Given specified values of the free (and fixed) parameters, in addition to a random seed value, networks are initialized, and numerical integration of the ordinary differential equations is entirely performed within `Brian2` (see Stimberg et al. [101], or online documentation for details). For all simulations, we use the forward Euler method with an integration time step of 0.2 ms, and networks are simulated for 200.1 seconds. The spike times of randomly selected 200 excitatory cells and 40 inhibitory cells are recorded throughout the simulation, and saved at completion.

To increase simulation efficiency, there is an early stopping criterion to detect networks that fail to generate activity or maintain the maximum possible firing rate. Simulations are paused after 10.1 seconds, where the average firing rate of all recorded excitatory neurons between 0.1 to 10.1 s is computed: 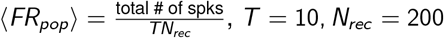. This population average firing rate is then compared to the maximum possible firing rate determined by the refractory period parameter of the excitatory population, i.e., *FR*_*max*_ = 1*/t*_*refrac*_. If ⟨*FR*_*pop*_⟩ *<* 0.0001 · *FR*_*max*_ or *>* 0.99 · *FR*_*max*_, the simulation is stopped, since the network will not produce meaningful activity, and the current parameter configuration is marked as “early-stopped”. Otherwise, the simulation is run to completion.

#### Simulation data organization and storage

Simulations are uniquely identified by two random seeds: batch and id. The batch seed is used for sampling parameter configurations, i.e., sample model configurations from the prior distribution or trained deep generative model. The id seed identifies configurations within the batch, and is used to sample stochastic components within a single network simulation, such as Poisson spikes and connectivity matrix.

Raw simulation data is composed of single-unit spike timestamps saved in hdf5 files and the corresponding parameter and seed values saved in csv files, along with various meta-information such as early-stopping status and simulation clock run-time. Various network activity summary features (described in next sections) are further extracted and saved in the same csv file. Given identical free parameter, batch, and id seed values, simulations can be reproduced exactly (up to numerical precision differences).

#### Prior predictives simulation dataset (i.e., training set)

To build the large database of simulations (e.g., visualized in Fig.2f and g), we sampled from the 28-dimensional uniform distribution 1,000,000 times. The support of this prior distribution is given by the 1-dimensional bounds defined for each free parameter in Table 1. Of these 1 million parameter configurations, 678,575 were early-stopped based on the above procedure. A further 58,230 failed subsequent quality assurance checks, where a simulation was discarded if: 1) the average single-unit ISI CV is less than 10^−7^, or 2) the population firing rate power spectrum contains 0 or infinity. This procedure further removed simulations that had constant near-zero or near-maximum firing rate, but was not caught in the early stopping check (see example PSDs in Supplemental Fig. S1d). The remaining 263,195 simulations were labeled as “valid” and submitted to further analyses for visualization or computing summary features as training set for the deep generative models.

### Network activity summary features

A variety of summary statistics were computed from network simulations, both for visualization and as target features during inference. The first 5.1 ms of simulations is discarded as burn-in time to allow networks to settle after initialization. Thus, all features are computed in the period of 5.1 to 200.1 seconds. Unless otherwise specified, features are computed based on spikes from all recorded excitatory neurons only. Analyses are performed in Python, largely relying on `numpy` [102],, `scipy` [103], and `scikit-learn` [104], while visualization is facilitated by `matplotlib` [105] and `seaborn` [106].

#### Single-unit ISI statistics

Single-unit features were computed based on inter-spike intervals (ISI), and averaged over excitatory and inhibitory neurons in the network separately. ISI distribution is defined as the set of intervals between two spikes, {*t*_*i*+1_−*t*_*i*_ }_*i*=1…*T*−1_, for all *T* spikes. Single-neuron ISI mean and standard deviation are computed accordingly, and the coefficient of variation (CV) is defined as *CV*_*ISI*_ = *σ*_*ISI*_ */µ*_*ISI*_. Finally, ⟨*µ*_*ISI*_ ⟩, ⟨*σ*_*ISI*_ ⟩, ⟨*CV*_*ISI*_ ⟩ are computed as means over recorded neurons in the network, which summarize a single network simulation (e.g., in Fig. 2f). Mean firing rate is computed as the inverse of the population average mean ISI. Neurons with fewer than 3 spikes are excluded.

#### Network dimensionality (PCA and participation ratio)

For a single simulation, single-unit firing rates were computed by binning individual spike trains of excitatory neurons at 10 ms resolution and then smoothed with a Gaussian window with standard deviation of 50 ms. The smoothed firing rate matrix is then subjected to Principal Component Analysis using `sklearn.decomposition.PCA` with 100 components. Total variance and percent variance explained are recorded (e.g., Fig.2e).

Participation ratio [66] is computed as the squared sum of eigenvalues divided by the sum of squared eigenvalues, and normalized by the number of PCs considered: 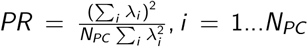, for various values of *N*_*PC*_. The normalization by *N*_*PC*_ ensures that the maximum dimensionality, regardless of the number of PCs considered, is 1.

#### Population firing rate

Single-unit spikes are binned at 1 ms resolution (i.e., 1000 Hz sampling rate) and averaged across recorded neurons of excitatory or inhibitory populations separately. For visualization and network burst extraction, the population rate time series is smoothed using a Gaussian window with 5 ms standard deviation.

#### Network bursts

First, the maximum of the smoothed population firing rate is found. Next, a small amount of Gaussian noise (*σ*=10^−7^) is added to the firing rate to avoid duplicate peak heights, and peak times are detected using `scipy.signal.find_peaks`, with the following arguments: `prominence=0.8`·`max_rate`, `distance=500` (i.e., 0.5 seconds), `wlen=20` seconds. Inter-burst interval (IBI) is defined analogous to ISI using burst peak times, while IBI mean and CV are similarly computed, per simulation. Given peak times, burst widths are found using `scipy.signal.peak_widths`, with `rel_height=0.95`. These features—IBI mean, IBI CV, and burst width—correspond to those shown in Fig. 3 and 4.

#### Population rate power spectral density (PSD)

PSD of the (unsmoothed) 1000 Hz population firing rate is estimated using Welch’s method (`scipy.signal.welch`), with default Hann window, 2-second segments and 1.5-second overlap (i.e., `nperseg=2000`, `noverlap=1500`), resulting in a frequency resolution of 0.5 Hz.

For the visualization in Fig. 2g, t-SNE is performed on *log*_10_*PSD* using `sklearn.manifold.TSNE` with the following parameters: `n_components=2`, `perplexity=30`, `learning_rate=200`, and `init=‘random’`. For computational tractability, 50,000 simulations were used instead of all 260k valid ones in the training database. Results are consistent using different random seeds and different subsets of simulations (Supplemental Fig. S2).

### Automated model inference from neural dynamics (AutoMIND)

In this section, we briefly describe the general methodology of simulation-based inference [50], as well as the specific variant we use, i.e., Neural Posterior Estimation [24] with Normalizing Flows [69]. Following the general descriptions, we detail the specific neural network architectures, training hyperparameters, and data pre-processing steps used in the various experiments. Finally, we describe the modified two-step sampling procedure specific to AutoMIND introduced in this paper. Training, inference, and sampling operations are facilitated by the open-source package `sbi` [68], which further utilizes `pytorch` [107] to perform neural network optimization.

#### Simulation-based inference (SBI) and Neural Posterior Estimation (NPE)

In SBI, a probabilistic deep generative model is optimized to perform conditional density estimation on mechanistic model parameter configurations and the corresponding model simulations. We refer to the *N*_*sims*_ pairs of parameters and simulations as 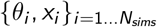. In this context, *θ*_*i*_ is always a 28-dimensional vector representing the values of the free parameters of the spiking neural network model in Table 1, while *x*_*i*_ is a k-dimensional vector representing the specific network activity features of interest, which we detail below. A simulations-only training dataset is generated by randomly sampling from a prior distribution over parameters, *θ* ∼ *p*(*θ*), and running them through the simulator model which defines an implicit likelihood, *x* ∼ *p*(*x* |*θ*). Here, we draw 1,000,000 independent samples from the 28-dimensional uniform prior distribution, and simulate spiking neural network activity as described above.

NPE aims to approximate the Bayesian posterior distribution for a specific target observation *x*_*target*_ with a deep generative model,

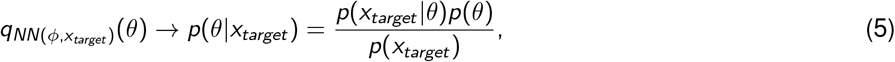

where:

- *q*(*θ*) is the approximating distribution, whose parameters are returned by flexible functions, such as artificial neural networks
- *NN*(*ϕ, x*) is an ANN with optimizable weights *ϕ*, and take as input *x*
- *x*_*target*_ is the target observation for which we want to infer candidate parameter configurations

With the training simulation dataset 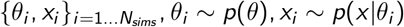, NPE optimizes the weights of the neural network by minimizing with respect to *ϕ* the loss function,

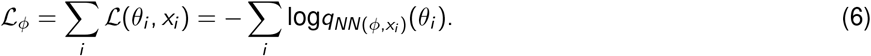

In other words, given features of simulated population activity *x*_*i*_, the neural network weights are optimized to return an approximating distribution *q*(*θ*) that maximizes the probability of the corresponding generating parameter configuration *θ*_*i*_, for all pairs of *θ*_*i*_, *x*_*i*_. It has been shown that by minimizing this loss function, the approximating distribution converges to the true posterior distribution [49].

Once trained, the deep generative model can be queried at inference time for posterior distributions given different target observations, including real experimental data, by passing into the learned neural network *x*_*target*_ and acquiring (samples from) 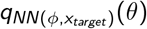. In particular, the “single-round” NPE variant used here shifts the cost of simulation, network training, and posterior sampling upfront, i.e., is amortized, such that at inference time, posterior samples can be acquired for any arbitrary *x*_*target*_ by simply sampling from the neural density estimator.

#### Normalizing Flow and Neural Spline Flow (NSF)

In all experiments, we use Normalizing Flows (NFs) as the deep generative model for conditional density estimation 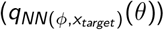 to approximate the posterior distribution. NFs are composed of a sequence of invertible transformations that morph a simple multivariate “base” distribution (e.g., standard Normal) into an arbitrarily shaped target distribution (i.e., the posterior distribution) of the same dimensionality. The transformations themselves are bijective functions that are easily invertible. The coefficients of the transformations, however, are input-dependent and parameterized by nonlinear neural networks, which at each stage takes the output of the previous stage as the input, in addition to the observation *x*_*i*_ as a conditioning variable in the context of conditional density estimation. Finally, an optional, simultaneously-trained “embedding network” can be prepended to first pre-process and reduce the dimensionality of the conditioning input *x*_*i*_, before passing the embedded data into each flow transformation block.

Specifically, we use Neural Spline Flows (NSFs) [70] with coupling layers. NSF transformations are composed of monotonic rational-quadratic splines with a user-specified number of segments (bins), and all spline coefficients are input-dependent and given by the output of artificial neural networks. The splines define how the base distribution is morphed along each segment of the support. See Papamakarios et al. [69] and Durkan et al. [70] for more details on NFs and NSFs, respectively.

#### Neural network architectures and training

In all experiments, we use NSFs with a stack of 5 transformations, each with 10 bins (spline segments). Spline coefficients are given by a residual neural network with 2 residual connection blocks, 50 hidden units each, and ReLU activation function. In addition, we use a fully connected multi-layer perceptron with 2-layers, 100 hidden units each, and 25 output units as the embedding network to pre-process the conditioning variable *x* before entering the flow transformations.

Neural network training uses `sbi` default settings: 90/10 train/validation set split of simulation dataset, batch size of 50, learning rate of 0.0005, and a stopping criterion of 20 epochs with non-decreasing validation loss.

#### Training data pre-processing and standardization

Two different sets of network activity features are computed on all fully completed simulations (321,425/1,000,000), as detailed above. In addition, prior to being used for training deep generative models, both datasets undergo a number of quality control checks and pre-processing steps to keep only high-quality simulations, such that the neural density estimator does not waste capacity in learning obviously unrealistic population dynamics. In both cases, the discarding thresholds were determined manually by examining model simulations, while log_10_-features were taken so that data features are more normally distributed and can be rescaled to be within good numerical ranges to facilitate better neural network training.

Burst timing features are composed of three population firing rate burst features: interburst interval (IBI) mean, CV, and burst width (i.e., three-dimensional vector). Simulations with less than two detected bursts, mean IBI *<* 0.01 seconds, IBI CV *<* 10^−4^, or burst width *<* 10^−4^ seconds are discarded. The log_10_ of these three features are taken, and each feature is standardized independently by subtracting the mean and dividing by the standard deviation over the training set.

PSD features are composed of the broadband power spectral density of the population firing rate from 0.5 to 495 Hz, at 0.5 Hz resolution (i.e., 990-dimensional vector). Simulations were further discarded if they satisfied any of the following conditions: 1) any frequencies had 0, inf, or NaNs, 2) average power between 2-10 Hz was below 10^−10^, 3) the variance of log_10_PSD *<* 7 × 10^−4^ or *>* 6.5, or 4) the range of log_10_PSD *<* 0.14 or *>* 13. Of the accepted samples, the log_10_ of PSD is taken, and standardized *per sample* by subtracting the mean across frequencies and divided by 10.

#### Posterior pseudo-mode sampling and error-based filtering

Given a target observation *x*_*target*_ at inference time, the trained Normalizing Flow can be directly queried to produce samples from the approximate posterior, 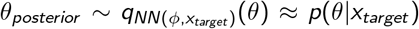. Due to unaccounted stochasticity within the simulator (e.g., instantiation of random connectivity, Poisson spikes), however, simulating these posterior samples with a particular random seed can produce posterior predictive simulations *x*_*predictive*_ that do not resemble the target observation. While these samples are nonetheless valid samples from the “correct” posterior distribution prescribed by Bayesian inference, in this study we instead prioritize the discovery of parameter configurations *confirmed* to be consistent with the target data, such that subsequent analyses are performed on ensembles of data-reproducing model configurations *only*. In doing so, we additionally bypass the potential concern that the approximate posterior coverage is overly broad. To accomplish this, we extend the sampling procedure in two ways:

First, we “oversample” the approximate posterior by an integer factor *k*_*oversample*_ and take the top *N*_*sample*_ samples with the highest log-likelihood under the posterior. For example, to acquire *N*_*sample*_ = 1000 samples in the end, we first sample *k*_*oversample*_ × 1000 parameter configurations from the approximate posterior, then evaluate the log-probability of those samples by querying the flow, and finally take the top 1000 samples with highest log-probability. This is equivalent to sampling around the modes (i.e., high-density regions) of the posterior—hence pseudo-mode sampling—and is efficiently done since both sampling and likelihood-evaluation are fast operations with Neural Spline Flows. Compared to naively sampling the posterior (i.e., *k*_*oversample*_ = 1), we find that pseudo-mode samples result in data-consistent predictive simulations much more consistently.

Second, we simulate all model configurations from the previous step and compute the target summary features accordingly (*x*_*predictive*_). We then compute the error between the target observation and all predictive simulations, i.e., _*D*_ |*x*_*target*_ − *x*_*predictive*_ |^2^, where *D* is the dimensionality of the summary feature (i.e., 3 for burst-timing and 990 for PSD). The final ensemble of candidate samples are taken as the *N*_*best*_ samples with the lowest errors.

For every target observation, we use *k*_*oversample*_ = 50 and *N*_*sample*_ = 1000 for inference on the synthetic dataset, and *k*_*oversample*_ = 100 for inference on both experimental datasets, with *N*_*sample*_ = 2500 for the organoid recordings and *N*_*sample*_ = 2000 for the mouse recordings. For visualization and analyses of discovered models, we use *N*_*best*_ = 100 or 200, as specified in the Results section. We emphasize that while, for example, *k*_*oversample*_ *N*_*sample*_ = 200000 samples are drawn from the Normalizing Flow for each organoid target recording, only 2000 are simulated to acquire 200 best-fitting models for further analyses, resulting in an effective yield of 10%.

### Datasets

Here we detail the acquisition and processing of experimental and synthetic datasets used as target observations for inference. Organoid and mouse datasets are previously published and downloaded from publicly available repositories. Data processing differ only in how the population firing rate time series is acquired from the raw spike train data, whereas network activity summary features are computed and standardized in the same way as for the simulation dataset, as described above.

#### Human induced pluripotent stem cell-derived brain organoid MEA recordings

Multi-electrode array recordings from human brain organoids were previously published in Trujillo et al. [51]. Preprocessed spike time data is accessed from the online repository [108] and a subset of recording days from one well (“Well 7”) were used in this study. Briefly, brain organoids were generated from human iPSC lines and maintained longitudinally for over 40 weeks. 2-3 organoids are plated on planar multi-electrode arrays with 64 low-impedance (0.04 MΩ) platinum microelectrodes with 30 *µ*m of diameter spaced by 200 *µ*m. Four-minute recordings were acquired around twice a week using a Maestro MEA System, 24 hours after each medium change, starting from 8-week post differentiation. Raw voltage traces are recorded at 12.5 kHz, and multi-unit activity from each channel were extracted by band-pass filtering between 300–3000 Hz and detected with an adaptive threshold of 5.5 standard deviations. Threshold crossings in each channel are registered as a MUA spike, which are binned at 1 ms resolution and averaged across 64 channels to form the population firing rate vector. Burst-timing features were then extracted as described above, while standardization at inference time used the mean and standard deviation of the training set. In the pharmacology experiment, a baseline recording was performed at week 27, followed by a perturbation recording 15 minutes after the addition of 10 *µ*M bicuculline.

#### Mouse cortical and hippocampal Neuropixels recordings

Mouse hippocampal and cortical recordings were acquired from the Allen Brain Observatory: Visual Coding Neuropixels Dataset [52], where mice were head-fixed and shown a variety of visual inputs while electrophysiological activity was recorded. Data pre-processing, quality control, and extraction of single-unit activity is described in the Dataset Release Technical Whitepaper. Spiketrain data was directly downloaded and accessed in Python using the `allensdk` package, version 2.15.2.

We use a single recording session, Session 771160300, with the acquisition timestamp 2018-11-05 13:14:59-08:00. In this session, over 900 single units are recorded across multiple locations, including sub-regions of visual cortex, thalamus, and hippocampus. We bin spike timestamps at 1 ms resolution while grouping units into one of 10 regions based on provided labels to compute the population mean firing rate. We selected two different blocks within the recording session as inference targets to minimize complex spatiotemporal structures in visual input: a 280 s block of continuous, stimulus-free spontaneous activity, and a block of trials with full-field dark/light flashes 250 ms each, separated by a 2 s inter-trial interval.

For the spontaneous activity block, PSD of the population firing rate was computed using Welch’s method as described for simulations above, with 2 s window and 1.5 s overlap. For the flash block, we separate each trial into two segments: “flash-on” between -0.25–0.75 s, and “flash-off” between 0.75–1.75 s, where 0 is relative to the onset of the flash. PSD was computed by averaging the squared-amplitude of Hann-windowed FFT (padded to 2000-samples) across flash-off or flash-on segments of all trials, resulting in a single PSD per area, per condition.

In total, this resulted in 10 regions×3 conditions=30 observations in the form of PSDs from this dataset, from which we select 16 different ones for inference. Similar to the simulation dataset, log_10_PSD was computed, and standardization is performed for each sample by subtracting its mean across frequencies and dividing by 10.

See Table 4 for detailed information about the target recordings, including number of units in each region and the selected conditions.

**Table 4.** Information on included brain regions and experimental conditions.

| Region | Full Name | Number of Neurons | Experimental Condition |
| --- | --- | --- | --- |
| CA1 | Cornu Ammonis 1 | 269 | Spontaneous, Flash Off |
| CA3 | Cornu Ammonis 3 | 42 | Spontaneous, Flash Off |
| DG | Dentate Gyrus | 29 | Spontaneous, Flash Off |
| ProS | Prosubiculum | 26 | Spontaneous, Flash Off |
| LP | Lateral Posterior Nucleus (Thalamus) | 43 | Spontaneous |
| VISp | Primary Visual Cortex | 85 | Spontaneous, Flash Off |
| VISam | Anteromedial Visual Cortex | 87 | Spontaneous, Flash Off |
| VISpm | Posteromedial Visual Cortex | 58 | Spontaneous, Flash Off, Flash On |

#### *In silico* circuit (synthetic) recordings

Synthetic targets were generated by simulating 1000 random parameter samples from the prior, which were excluded from the training dataset. 25 target observations were manually chosen based on the population firing time series and PSD, in order to include a wide variety of dynamics. Data pre-processing, summary feature extraction, and standardization before inference are performed in identical fashion as for the training set.

### Analysis, visualization, and perturbation of discovered models

All subsequent analyses and visualization are performed on the *N*_*best*_ candidate model parameter configurations and/or the corresponding predictive simulations.

#### Violin plots, kernel density estimation (KDE), and other visualizations

1 -dimensional smoothed histograms are generated using either `matplotlib.violinplot`, with internally computed KDE using `bw=0.25`, or `scipy.stats.gaussian_kde` with default argument values. Ticks denote median value, and whenever applicable, 25th and 75th percentiles. 2-dimensional KDE plots are generated using `seaborn.kdeplot`, typically with `levels=5` and `thresh=0.25`. To avoid visual clutter, raster plots of simulation spike trains are typically shown with every *N*_*skip*_ spikes plotted, where *N*_*skip*_ is either set to be 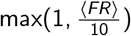 (in Fig. 2), or determined manually. Predictive simulation PSD plots show mean ± standard deviation of log_10_PSD.

#### Statistics

In all correlation analyses and 2-dimensional scatterplots, we report Spearman’s *ρ* and p-values computed using `scipy.stats.spearmanr`, where */**/*** denote *p* = 0.05*/*0.01*/*0.001, respectively (unless specified otherwise).

#### Perturbation experiments

In the organoid perturbation experiment (Fig. 4), GABA blockade was emulated in the discovered candidate models by setting all inhibitory synaptic strength parameters (*Q*_*I*→*E*_ and *Q*_*I*→*I*_) to 0 while keeping all other parameters identical.

In the mouse circuit input experiment (Fig. 5h), all candidate models were re-simulated with the Poisson input rate for excitatory populations (*ν*_*input*→*E*_) set to 150 % the inferred value. Perturb/baseline spectral power ratio was computed by taking the difference of log_10_PSDs, sorted according to the frequency of maximum power in the baseline models.

#### Parameter-shuffle analyses

To destroy higher-order relations between parameters (i.e., pairwise correlation and beyond) while maintaining the 1-dimensional marginal distributions, values of the *N*_*best*_ discovered models were shuffled for each parameter independently to create a new set of candidate models.

#### Subspace and sensitivity analyses

Principal component analysis was performed on 500 candidate models for each of the 25 synthetic target observations (“per-xo”). Then all 25 × 500 models were concatenated and randomly distributed into 25 groups, where PCA was again performed per group “mixed”. In both cases, we extract percent variance explained for each of the 25 groups, and compute the mean and standard deviation. For “mixed-xo”, we compute the percent variance explained when projecting the 500 samples from the same target observation onto the principal components estimated from the randomly mixed groups.

For the sensitivity analyses, we take each of the *N*_*best*_ models of a single target observation and conditionally sample that approximate posterior 100 times for a single parameter while holding fixed (i.e., conditioning on) the values of the 27 other parameters. This was repeated for all 28 parameters for a single discovered model, and for all *N*_*best*_ discovered models.

## Supplemental Materials

**Figure S1.**
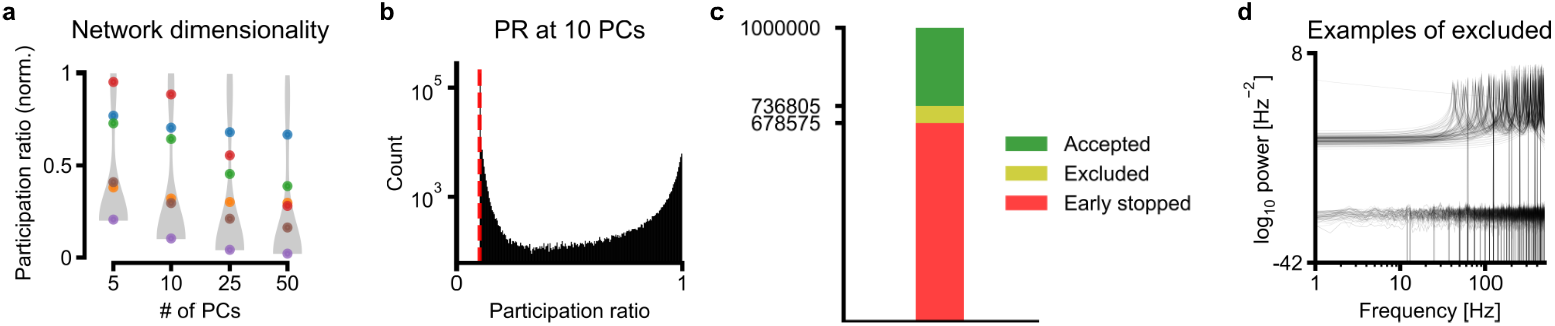
Additional statistics on the dataset of 1-million training simulations. **a**, Participation ratio computed with various numbers of principal components included. Violin denotes distribution over whole dataset, colored dots correspond to examples in Fig. 2c. **b**, Distribution of participation ratio over entire dataset, at 10 principal components. Red dashed line denotes minimum possible value. **c**, Numbers of early-stopped, excluded, and accepted simulations from 1-million parameter configurations. **d**, Power spectral density of example simulations that were completed, but excluded post-hoc based on quality control criteria (detail in Methods section).

**Figure S2.**
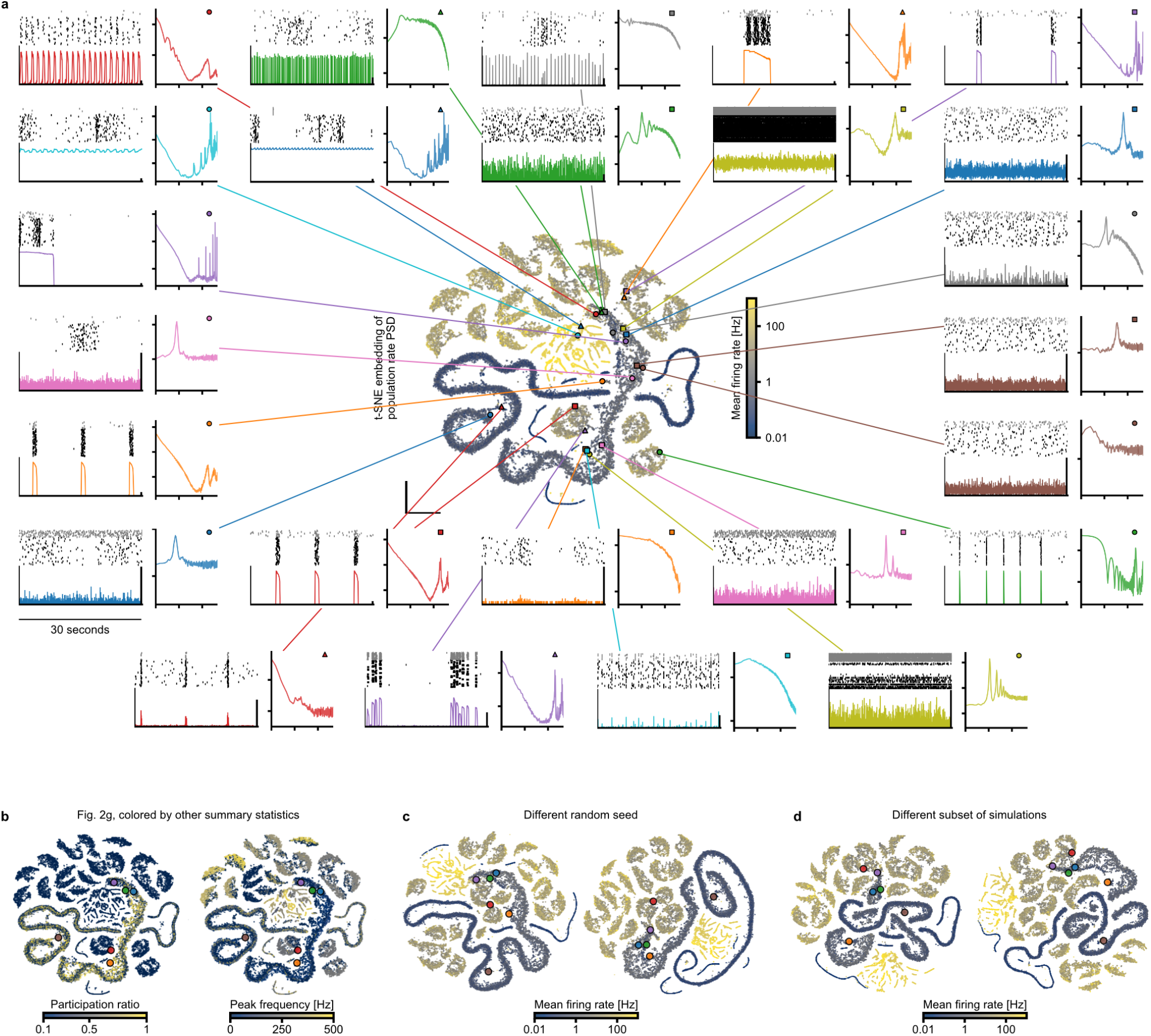
Additional examples of network simulations in t-SNE embedding. **a**, Visualization of additional simulations, including population raster, firing rate, and PSD, and their location in 2-dimensional t-SNE embedding. **b**, Same embedding as Fig. 2g, colored by non-spectral power features, i.e., participation ratio at 10 PCs and peak frequency of population rate PSD. **c**, t-SNE embedding on the same subset of 25,000 simulations as Fig. 2g, with two different random seeds producing qualitatively similar structures. **d**, t-SNE embedding with two different subsets of 25,000 simulations from the training dataset. All t-SNE embeddings are performed on broadband PSD of population firing rate.

**Figure S3.**
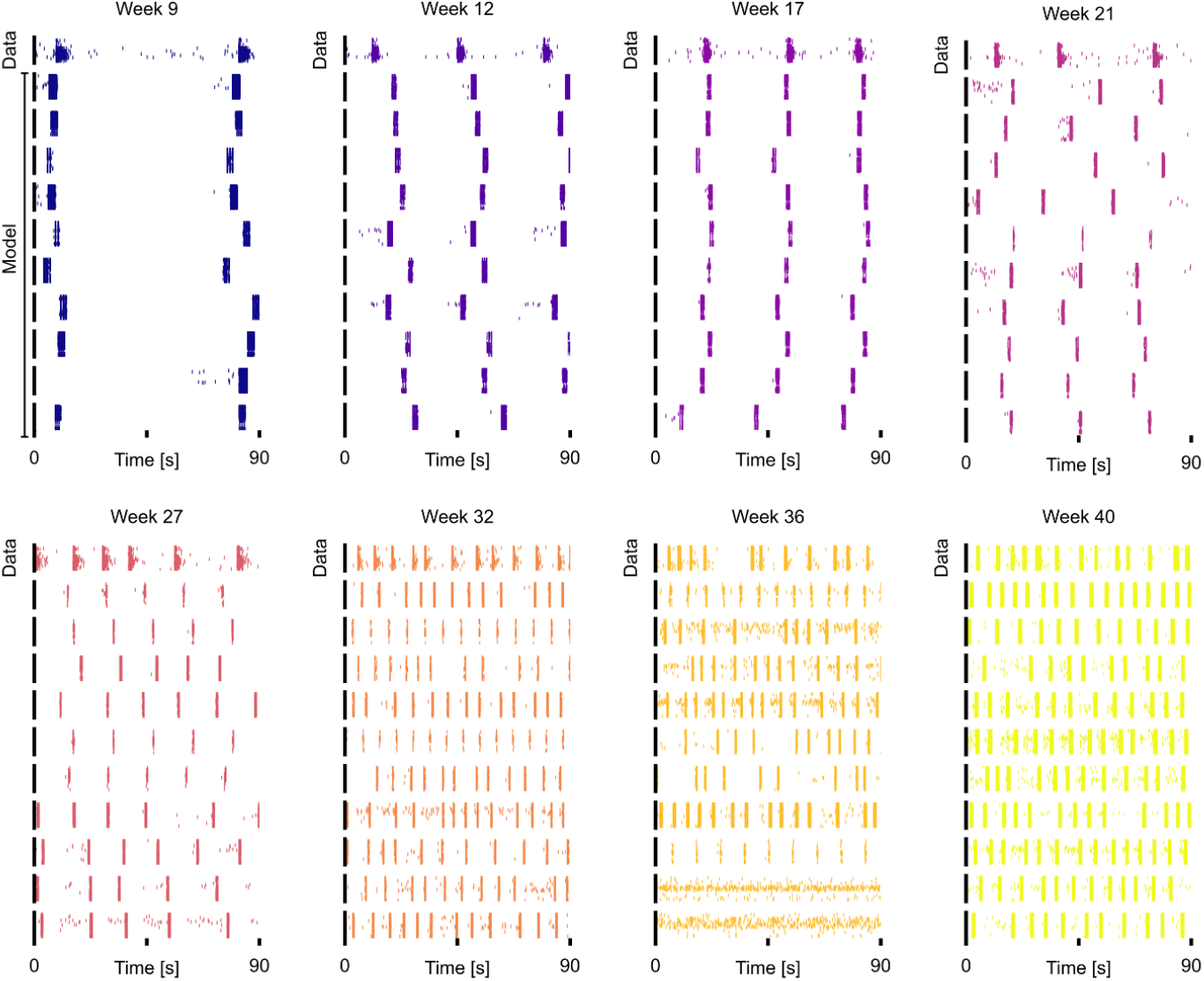
Additional discovered organoid model simulations. In each sub-panel, the first row shows real organoid recordings, i.e., target observations, while the rows below show 10 different examples of discovered models with burst summary features consistent with that recording.

**Figure S4.**
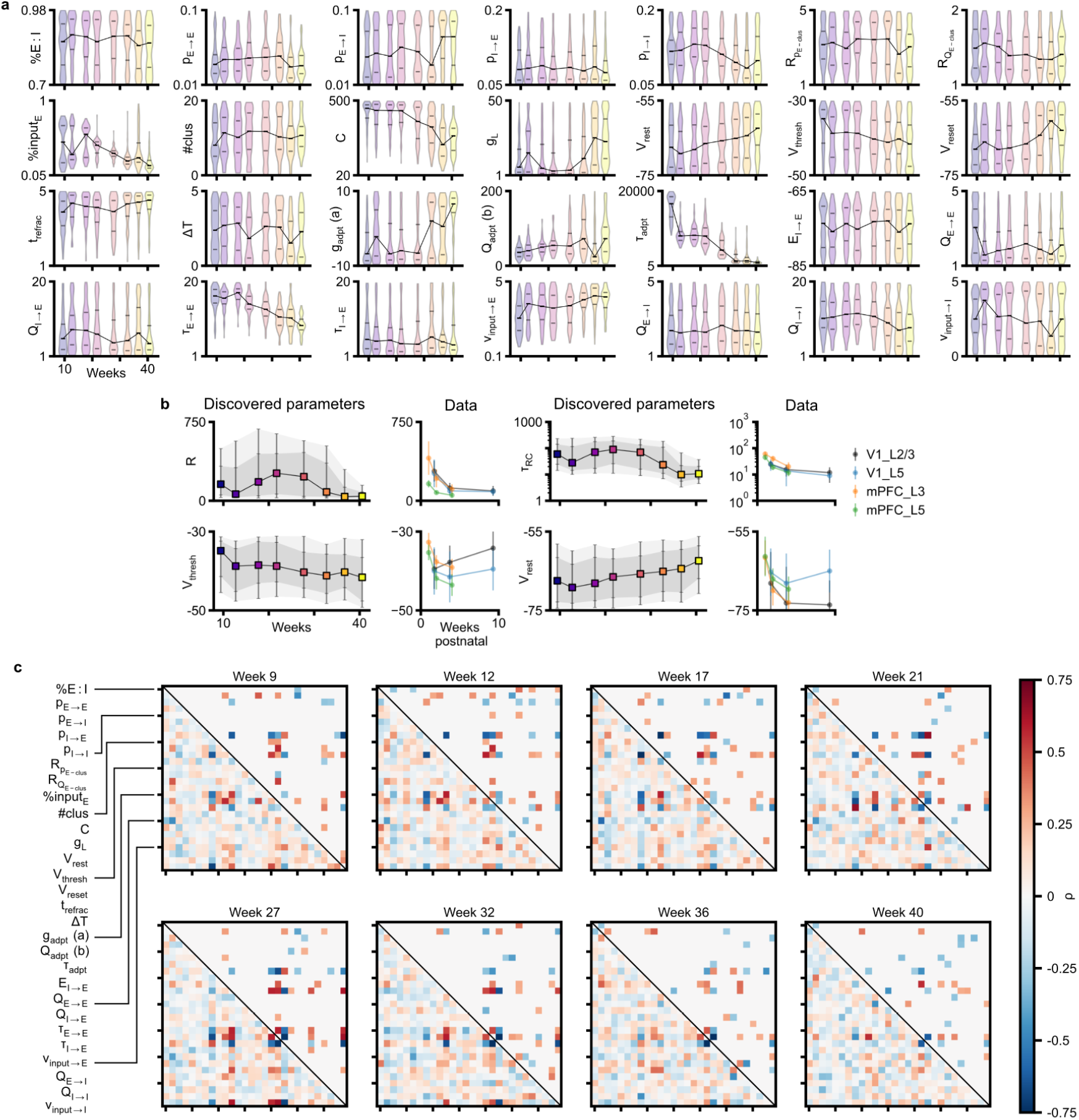
Discovered parameter configurations of organoid recordings. **a**, Evolution of all 28 parameters in discovered models over organoid development. Ticks on violin show 25th, 50th, and 75th percentile. **b**, Parameter predictions of discovered models compared to experimental values measured in early postnatal rodent cortex (See Main Text for references). **c**, Parameter correlations computed from ensemble of discovered models for each organoid recording (Spearman correlation). The lower triangular shows all correlations, and the upper triangular shows only significant correlations with *p <* 0.01.

**Figure S5.**
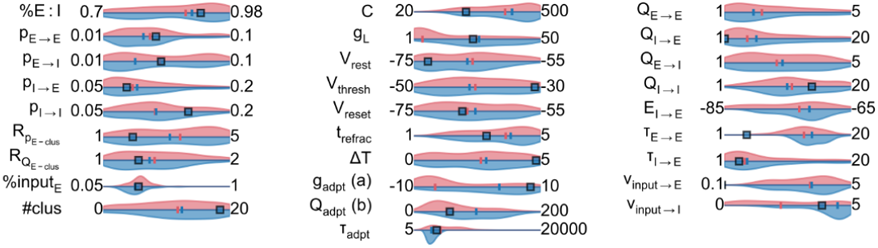
Parameter distributions of perturbation-consistent organoid models. Pink densities are computed over all discovered models of the baseline recording at week 27. Blue densities show only the subset of those models further consistent with the GABA-blockade perturbation recording after removal of inhibitory currents. Subset shown in Fig. 4d.

**Figure S6.**
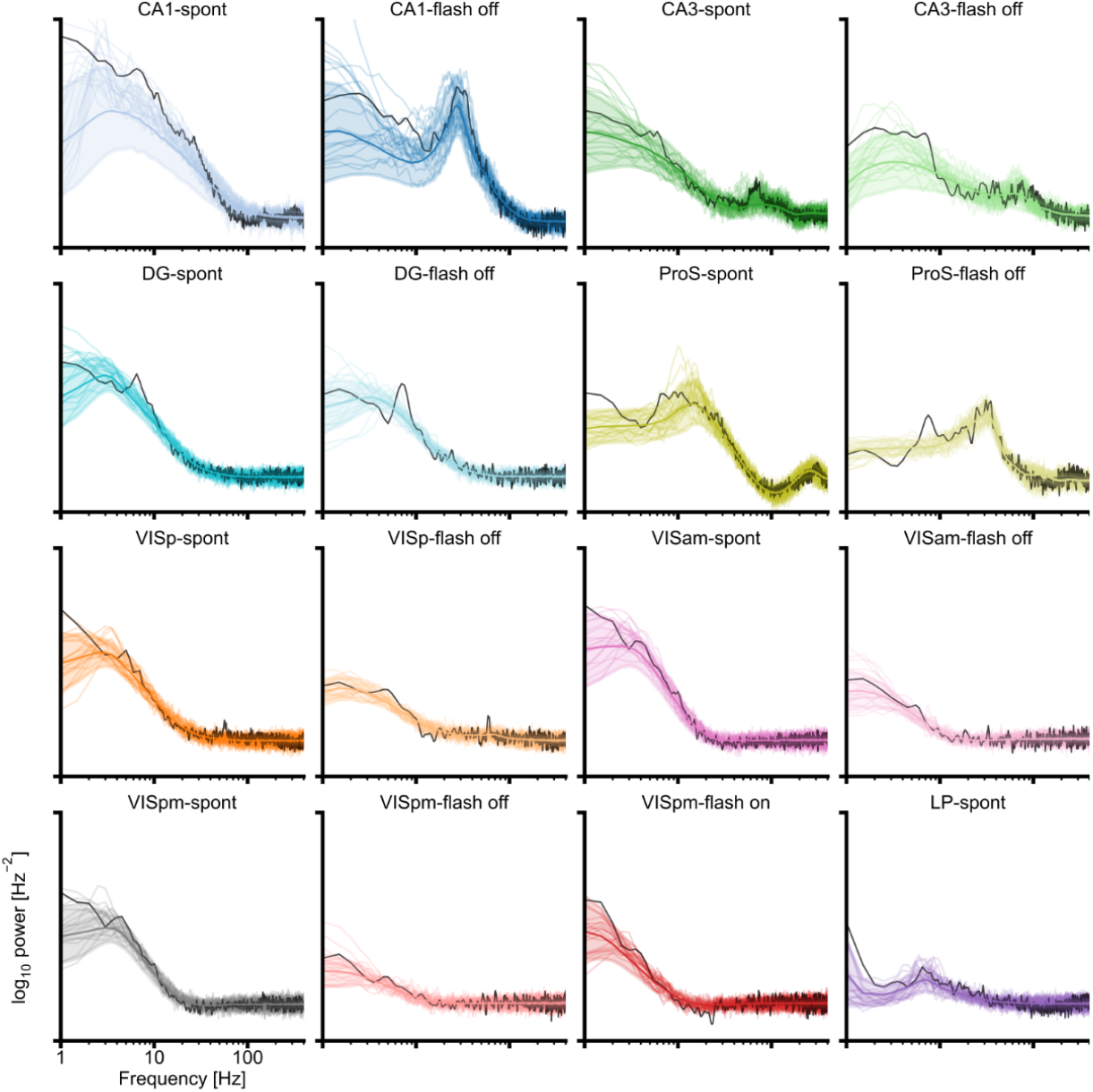
Discovered model simulations for all target observations in the mouse Neuropixels dataset. Black lines show PSD of target observations, thick colored lines and shading show mean and standard deviation over all discovered model simulations, and thin lines show 20 individual example models. Subset shown in Fig. 5b.

**Figure S7.**
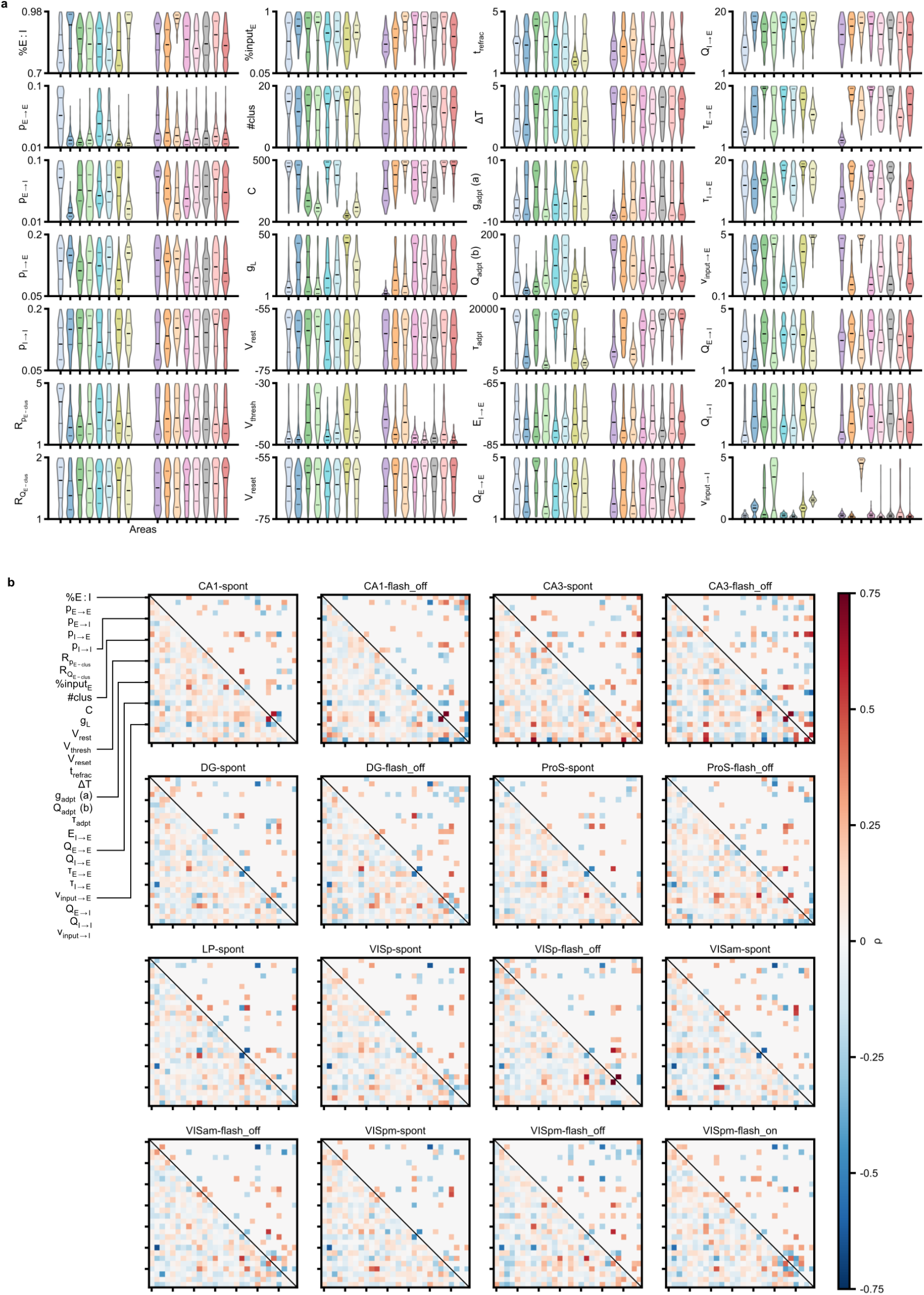
Discovered parameter configurations of mouse Neuropixels recordings. **a**, All parameters in discovered models of mouse hippocampus, thalamus, and visual cortex. Ticks show 25th, 50th, and 75th percentile. **b**, Spearman correlations; the lower triangular shows all correlations, and the upper triangular shows only significant correlations with *p <* 0.01.

**Figure S8.**
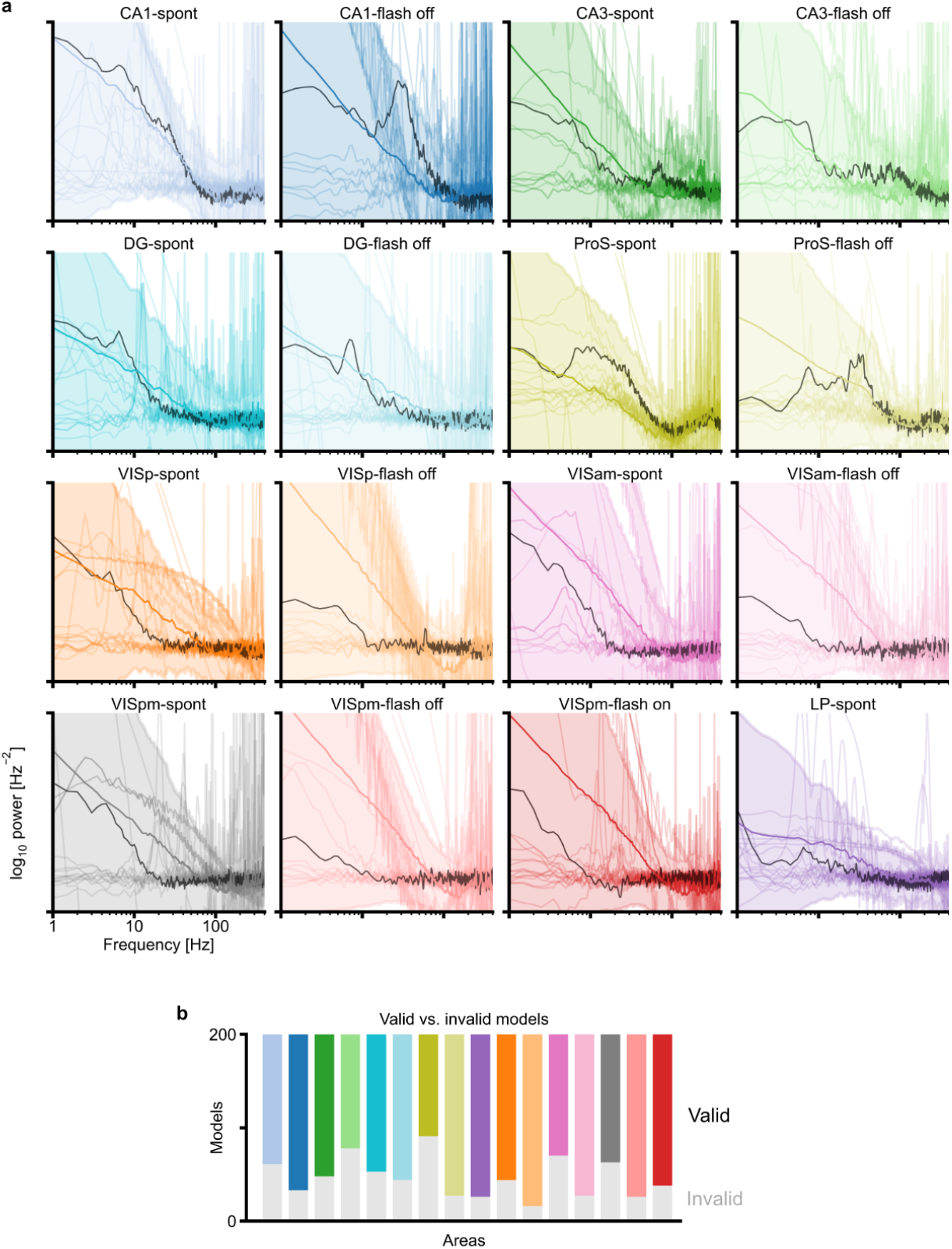
Simulations of shuffled models for mouse Neuropixels recordings. **a**, Population firing rate PSD of shuffled models. Black lines show PSD of target observations, thick colored lines and shading show mean and standard deviation over all shuffled model simulations, and thin lines show individual example models. Subset shown in Fig. 5h. **b**, Proportion of shuffled models that produced valid network simulations, based on the same exclusion criteria (e.g., Fig. S1d).

**Figure S9.**
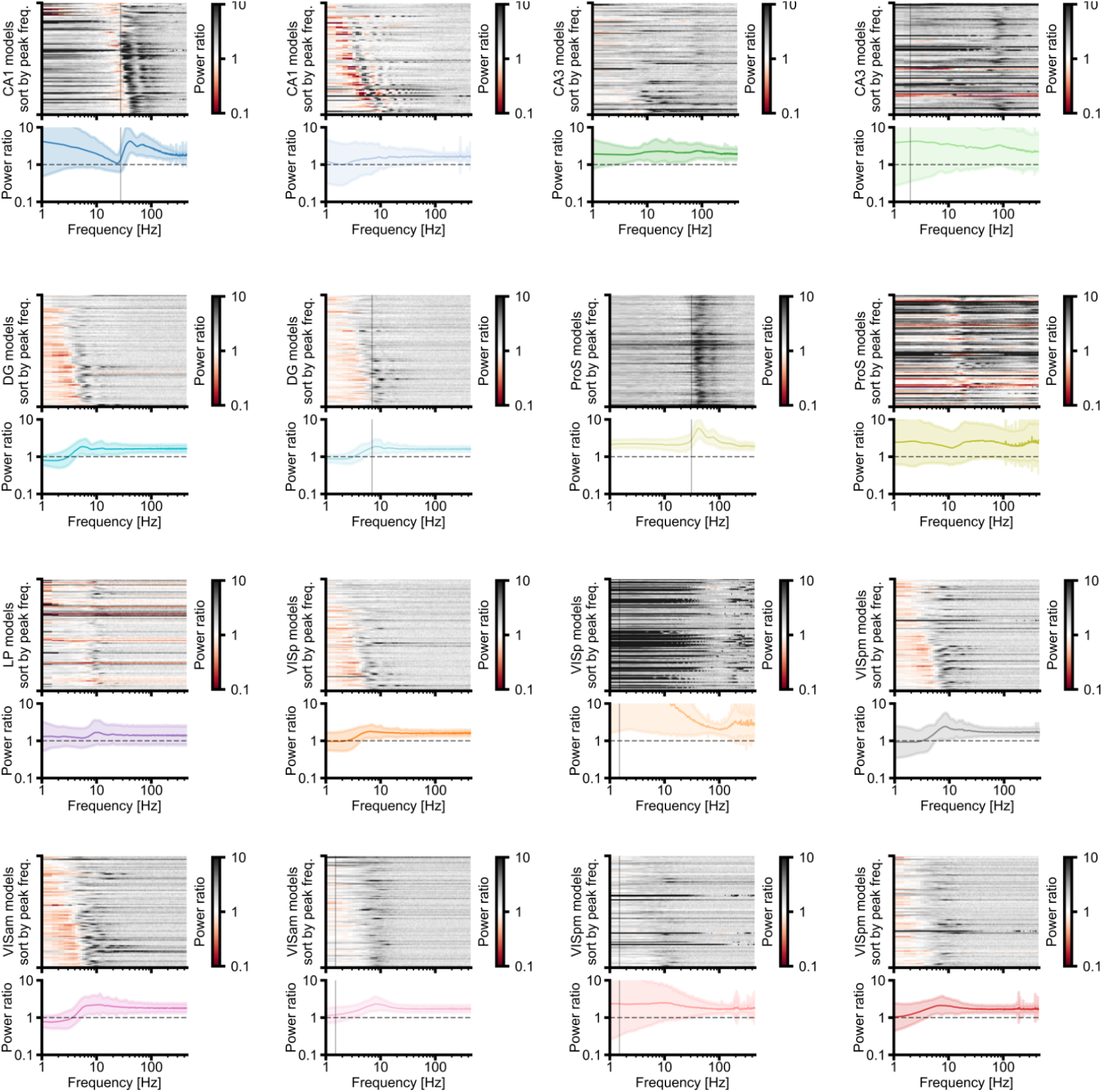
Response of all discovered models for mouse Neuropixels recordings after 50% increase in excitatory drive. The top panel of each sub-plot shows the responses (post-to-pre power ratio) of individual discovered models, sorted by increasing peak frequency; the bottom panel shows average and standard deviation of spectral response over all models. Vertical lines, when present, denote frequency of detected narrowband oscillation in the PSD of target observations. Subset shown in Fig. 5j.

**Figure S10.**
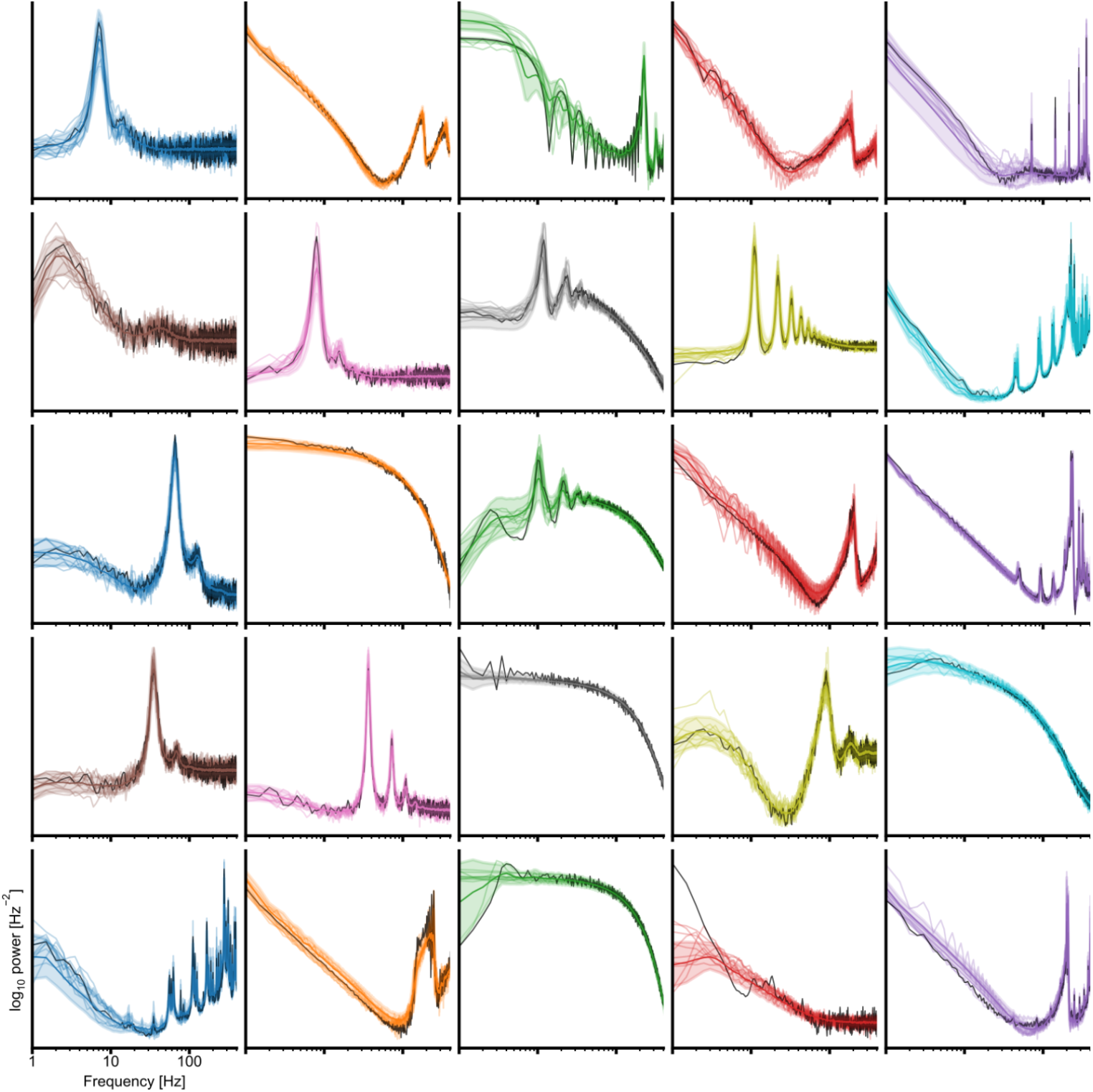
Discovered model simulations for all target observations in the synthetic dataset. Black lines show PSD of target observations, thick colored lines and shading show mean and standard deviation over all discovered model simulations, and thin lines show 20 individual example models. Subset shown in Fig. 6b.

**Figure S11.**
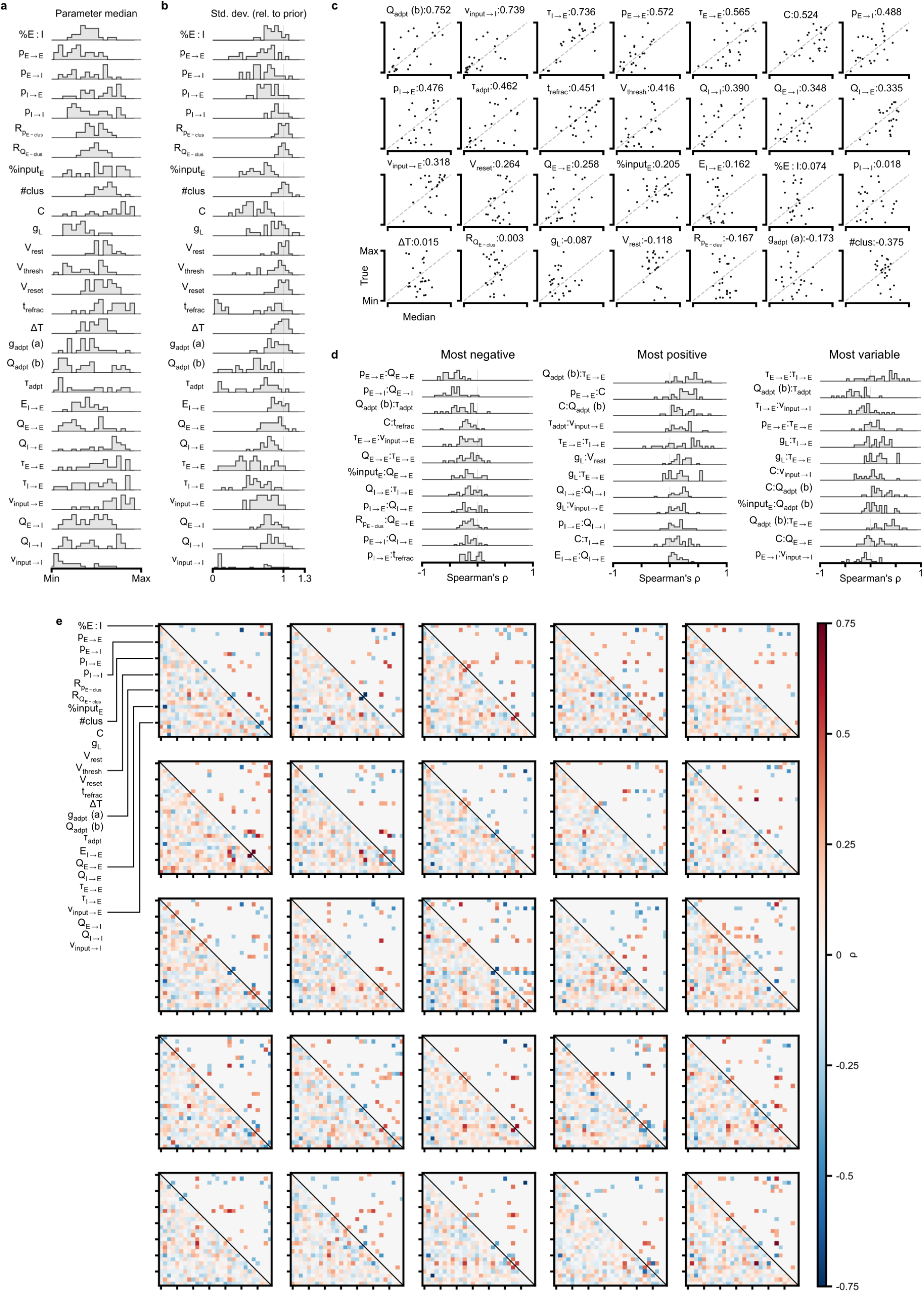
Discovered parameter configurations for all target observations in the synthetic dataset. a-d show aggregated statistics over ensembles of discovered models for all 25 target observations. **a**, Distributions of median value for all parameters. **b**, Distributions of standard deviation for all parameters. **c**, Parameter median vs. “True” value used in generating the target observation. **d**, Distributions of parameter correlations for a subset of parameter pairs, including those with the most negative (left), most positive (middle), and most variable (right) correlations. **e**, Spearman correlations over ensembles of discovered models for all 25 target observations; the lower triangular shows all correlations, and the upper triangular shows only significant correlations with *p <* 0.01.

**Figure S12.**
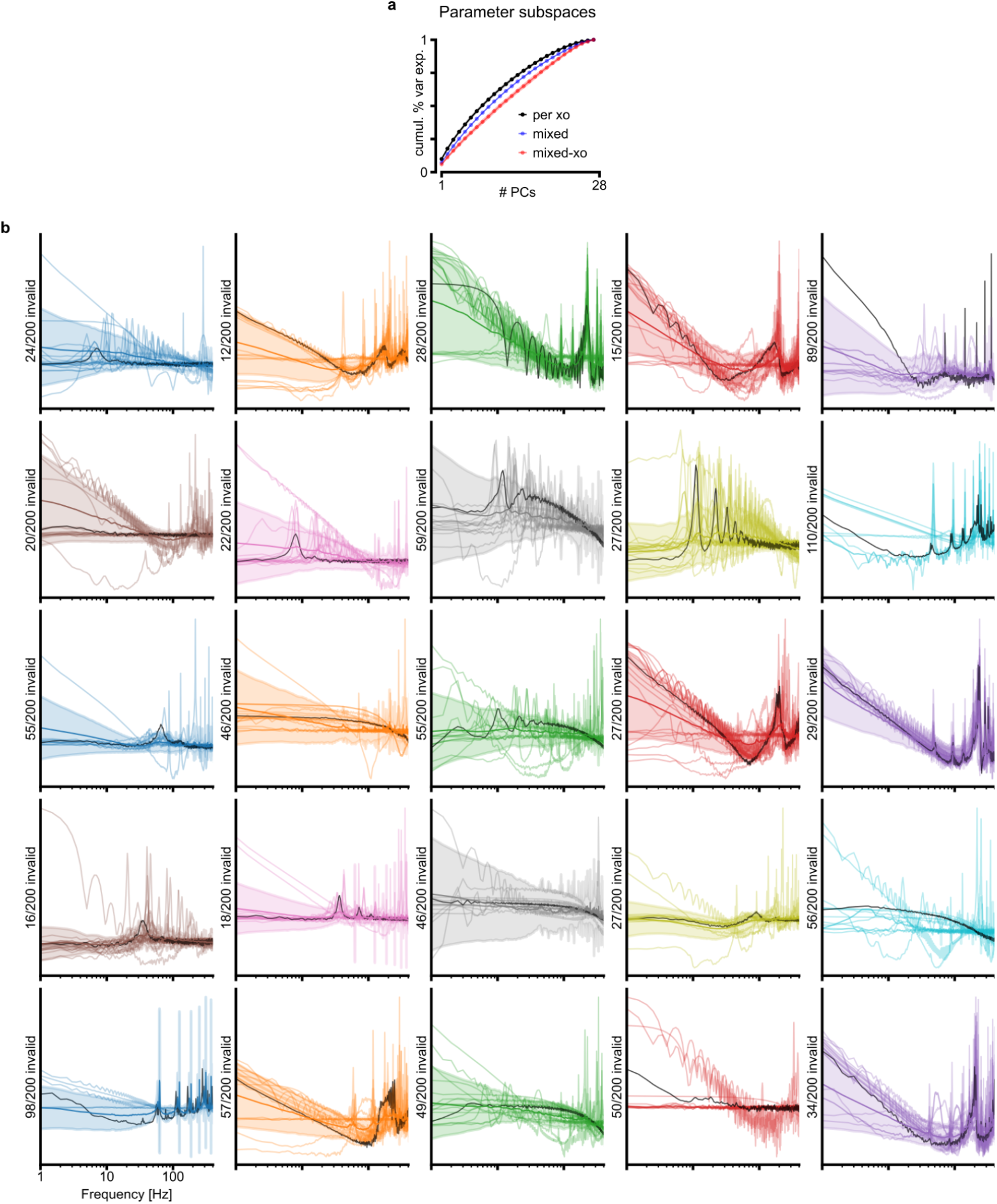
Parameter subspace analysis on discovered models of the synthetic dataset. **a**, per xo (black): cumulative proportion of variance explained when principal components are fit to the ensemble of 400 discovered models for each target observation; mixed (blue): variance explained when PCs are fit to random subsets of 400 models sampled across ensembles for different observations; mixed-xo (red): variance explained when projecting models of the same target observation (within-ensemble) onto PCs fit to randomly sampled subsets (across observations). Dots and shading show mean and standard deviation over 25 target observations / random subsets. **b**, Population firing rate PSD of shuffled models (i.e., parameter correlations destroyed). Black lines show PSD of target observation, thick colored lines and shading show mean and standard deviation over all shuffled model simulations, and thin lines show individual example models. “N/200 invalid” denotes the number of shuffled models whose simulations were early-stopped or rejected.

**Figure S13.**
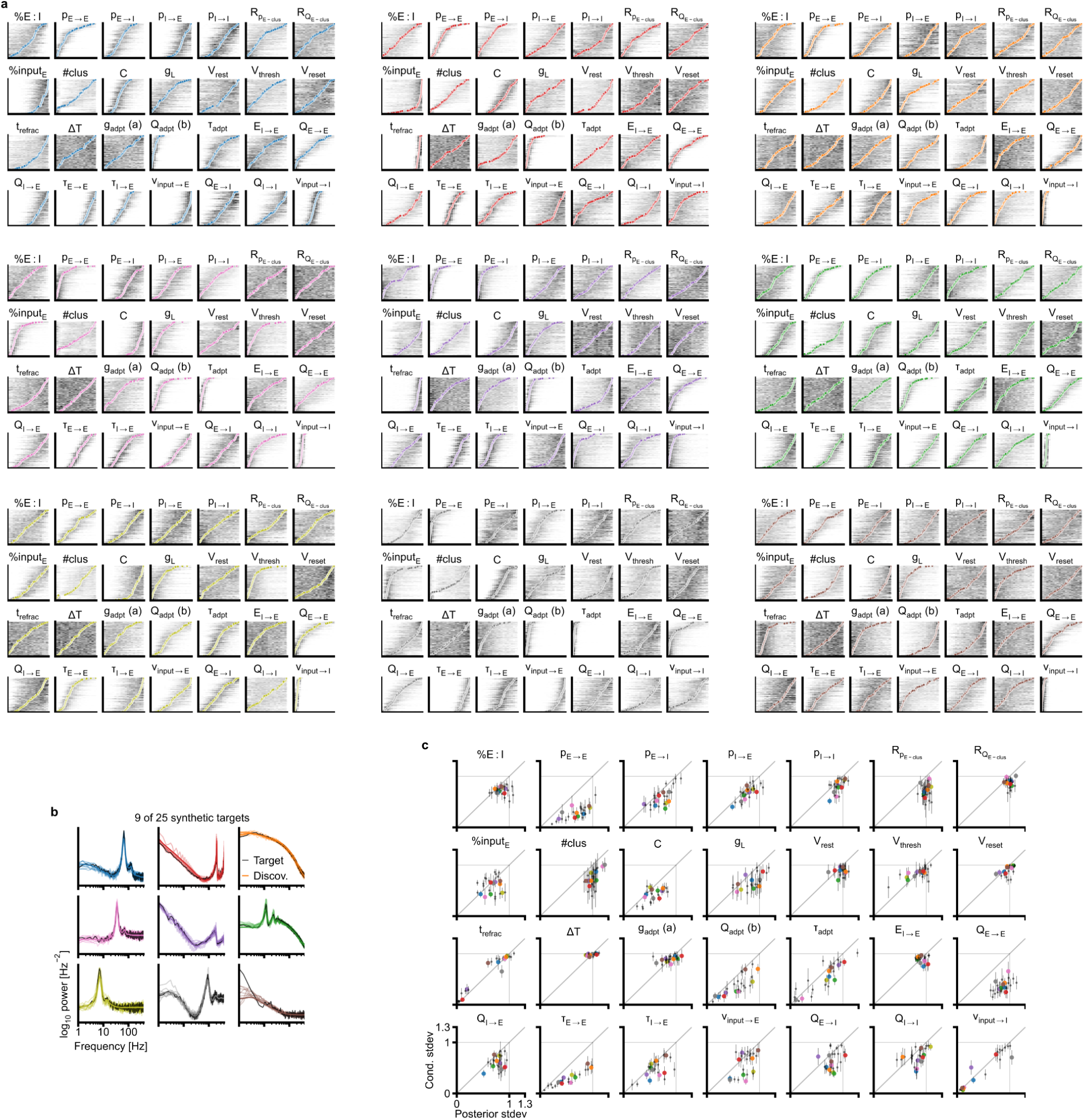
Local sensitivity analysis for all parameters and discovered models across target observations in the synthetic dataset. **a**, Small panels within each 7-by-4 grid displays the conditional density of one parameter (darker means higher density) when all other parameters are held fixed to the value of a single discovered model (colored dot), across all discovered models. **b**, The 9 example target observations and discovered model simulations corresponding to the 9 sub-panels in a. Same as Fig. 6b. **c**, Standard deviation of discovered model parameter values (Posterior stdev, same as Fig. 6f) vs. the mean standard deviation of conditional densities (Cond. stdev), across all discovered models for all target observations. Values are normalized by the standard deviation of the uniform prior distribution. Error bars show standard deviation of conditional standard deviation; colored dots correspond to the example target observations in b.

